# Metabolic and neuroactivity imbalances in plasma from aniridia patients with *PAX6* haploinsufficiency

**DOI:** 10.1101/2024.11.07.622475

**Authors:** Dulce Lima Cunha, Vivienne Kit, Jane Skinner, Ailsa A Welch, Mariya Moosajee

**Affiliations:** UCL Institute of Ophthalmology, Dept Brain Sciences, London, UK; Moorfields Eye Hospital NHS Foundation Trust, London, UK; Centre for Population Health Research, Faculty of Health, University of East Anglia, Norwich, UK; The Francis Crick Institute, London, UK; Great Ormond Street Hospital for Children NHS Foundation Trust, London, UK

## Abstract

PAX6 is a transcription factor crucial for the development of the eye, pancreas, and brain. Heterozygous variants resulting in *PAX6* haploinsufficiency are the main genetic cause of congenital aniridia, characterized by both anterior and posterior ocular defects and sight loss. The extra-ocular features of *PAX6* haploinsufficiency are becoming more widely recognised, with systemic manifestations like obesity, diabetes, and neurological/behavioural disorders being reported. In this study, we uncovered the metabolomic profile of the blood plasma from 25 *PAX6*-related aniridia patients compared to gender and age-matched controls. We found significant disruptions in lipid and energy metabolism, increased oxidative stress and neurotransmitters imbalances, as well as alterations linked to the gut microbiome. This study identified novel metabolic changes associated with *PAX6* haploinsufficiency, providing evidence for the systemic aetiology of congenital aniridia and emphasizing the need for multidisciplinary management and further exploration into ocular and systemic therapeutic approaches.

## Introduction

The paired box gene 6, *PAX6*, encodes a transcription factor essential for the development of the eye, pancreas and central nervous system (CNS). Biallelic variants in *PAX6* (OMIM #607108) usually lead to embryonic death but pathogenic heterozygous variants predominantly result in loss of the mutated transcript, leading to *PAX6* haploinsufficiency [1]. *PAX6* haploinsufficiency is the underlying mechanism behind the pan-ocular disease aniridia (OMIM # 106210), typically characterised by congenital iris hypoplasia, foveal hypoplasia and nystagmus, with later onset of cataracts, glaucoma and corneal opacities, resulting in significant vision loss [2]. Aniridia can occur in isolation or as part of the Wilms tumour, Aniridia, Genitourinary abnormalities and mental Retardation (WAGR) syndrome (OMIM # 194072), which results from larger deletions encompassing *PAX6* and neighbour gene *WT1* [3]. A form of WAGR syndrome that includes obesity (named WAGRO, OMIM # 612469) has been described in patients where deletions also encompass the brain-derived neurotrophic factor gene, *BDNF*, involved in energy homeostasis [4]. *BDNF* haploinsufficiency correlates with increased appetite and higher Body Mass Index (BMI) in both mice and humans [5, 6].

Ocular phenotypes in aniridia patients, albeit highly variable, have been thoroughly described [7–10]; systemic manifestations however, have been poorly characterised, despite growing evidence revealing the majority of patients also present with neuronal and/or endocrine abnormalities [11–16]. PAX6 plays a vital role in pancreatic development, being necessary for the formation of islets cell types [17]. Accordingly, mice carrying islet-specific Pax6 inactivation are born with reduced number of α, β, and δ cells and die soon after birth with severe diabetic phenotype [18–20]. Post-natally, Pax6 acts on maintaining the identity and function of mature islet cells and lack of expression results in a severe decrease in insulin, glucagon, and somatostatin production [21–23]. Metabolic issues reported in aniridia patients include obesity, hyperglycemia, insulin resistance and diabetes [13–16, 24–26]. Several studies have shown that patients have increased proinsulin levels and altered proinsulin/insulin and proinsulin/ C-peptide ratios [16, 27]. These observations were recapitulated in mice with the heterozygous *Pax6* p.(Arg266*) stop mutation, which showed altered proinsulin processing and defect in glucose metabolism due to prohormone convertase 1/3 deficiency [27].

Pax6 has a crucial role in establishing early brain territories [28]. It is expressed in both embryonic and adult neural stem cells, contributing to neurogenesis in the postnatal brain, where it is also expressed in the olfactory bulb, cerebellum and amygdala [28–30]. Structural abnormalities found in patients include pineal gland hypoplasia, absence or abnormalities in olfactory gland, anterior commissure or corpus callosum (reviewed in [29]). Consequently, patients can suffer from sleep disturbances and circadian rhythm alterations, neurodevelopmental disorders like autism or ADHD, depression, anxiety or auditory processing disorder [11, 30–32].

Similar to ocular phenotypes, there is a high variability of systemic manifestations seen in aniridia patients, with no clear genotype-phenotype correlations established. Also, no preventive treatments are available due to the lack of comprehensive analysis of these manifestations. Uncovering biomarkers for the identification of early stage metabolic abnormalities would therefore be fundamental to understand phenotypic differences and manage disease comorbidities.

Metabolomic analysis of human plasma has increasingly been used to elucidate disease mechanisms and allow the identification of novel biomarkers on a growing number of ocular diseases [33–37]. The aim of this study is therefore to characterise the metabolomic profile of aniridia patients and uncover the systemic consequences of *PAX6* haploinsufficiency. We identified strong signals from metabolites associated with oxidative stress, lipid metabolism, neuroactivity and links to the microbiome. Our findings reinforce the aetiology of aniridia as a systemic disorder, propose novel targets and mechanisms linked to *PAX6* haploinsufficiency and explore possible therapeutic approaches.

## Results

### 1. Cohort description

Twenty-five patients from 18 independent families clinically and molecularly diagnosed with aniridia caused by pathogenic *PAX6* variants were included in this study, together with 25 age and gender-matched unaffected individuals. Of these, 13 (52%) were female and 12 (48%) were male in each group. Age at the time of blood collection ranged from 8 to 64 years; the mean age was 31.88 ±16.71 for the aniridia group and 31.56 ±16.52 for the control group, comprising 6 (24%) children (under 18) and 19 (76%) adults. Mean of BMI values in the aniridia group was 27.7 ±6.1, which was significantly increased (Mann-Whitney test) compared to the control unaffected group (23.99 ±5.75, *p*=0.033). The ethnicity of the *PAX6* patient cohort consisted of White British (23/25), Black British (1/25) and Latvian (1/25).

Genetic and phenotypic characteristics are described in detail in Supplementary Table 1. The majority of patients (23 out of 25) included were diagnosed with likely pathogenic or pathogenic variants predicted to cause *PAX6* haploinsufficiency: 11 frameshift, 10 nonsense, 1 synonymous/predicted splicing variants and 1 with whole gene deletion. Two aniridia patients with variants leading to C-terminal extension (CTE) of the PAX6 protein were also included. The most common *PAX6* variant in the aniridia cohort was the nonsense c.718C>T, p.(Arg240*) in exon 9, present in 7 out of the 25 patients (28%).

Clinical evaluation showed high inter- and intra-familial phenotypic variability with no obvious genotype-phenotype correlations identified. All patients presented with a spectrum of iris defects from iris pigmentation and iris atrophy to partial and complete iris hypoplasia, with 23/25 presenting with different grades of bilateral foveal hypoplasia and 21/25 with nystagmus. Severity of secondary manifestations appeared to correlate with age and included presence of glaucoma (8/25), lens opacities/cataracts (19/25) and aniridia-related keratopathy (22/25). Interestingly, one patient was also diagnosed with chronic progressive external ophthalmoplegia.

Extraocular phenotypes were documented in all but 5 patients, but no correlation to the mutation type was seen. Obesity (BMI ≥30) was reported in 11/25. Gastrointestinal issues were also reported including gastric reflux in 2/25, irritable bowel syndrome in 2/25 and constipation in 1/25. Neurological issues such as anxiety (3/25), headaches and migraines (4/25), central auditory processing disorders (3/25), depression (2/25), seizures including functional seizures (2/25) and sleep disturbances (1/25) were also described. Structural abnormalities of the brain were present in 2/25. Learning difficulties were identified in three children, while one child was diagnosed with autism and two other were diagnosed with ADHD. Other systemic features reported included fibromyalgia (2/25), abdominal hernias (2/25), hypertension (2/25), eczema (1/25) and hypothyroidism (1/25). A comprehensive summary of all extra-ocular features are included in supplementary Table 1. Aniridia patients taking statins, those with type 1 or 2 diabetes mellitus, and patients with contiguous *PAX6* and *WT1* gene deletions causing WAGR syndrome were not included in this study.

### 2. Overview of metabolite and dietary differences in aniridia patients

Mann-Whitney tests were performed to evaluate dietary preferences of both groups following food frequency questionnaire [35, 38, 39], and no significant differences were found between the two groups regarding intake of any nutrient class (Supplementary Table 2). Principal component analysis (PCA) revealed little separation regarding overall metabolomic profiles between aniridia patients and their age-matched controls (Figure 1A), also found through global heatmap analysis (Supplementary Figure 1). However, some stratification by age was visible (Figure 1B).

**Figure 1.**
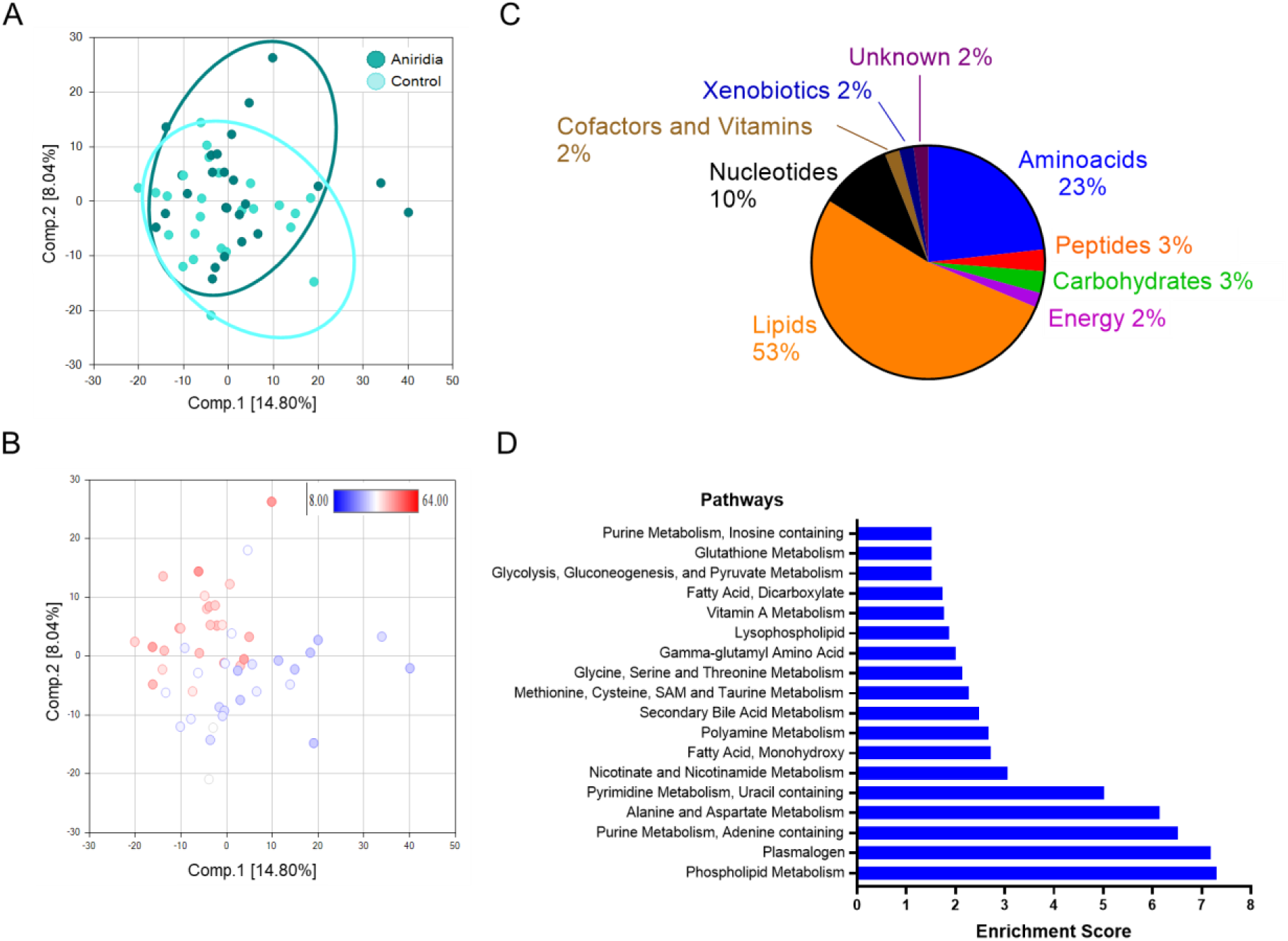
Global metabolomic analysis of aniridia patients versus age- and gender-matched controls. A) Principal component analysis (PCA) showing control and aniridia samples in light and dark blue, respectively (n=25 each group). B) PCA showing all samples distributed by age (8 to 64 years old). C) Distribution of significantly altered metabolites within their classes. D) MetaboLync pathway analysis software determined the most significantly altered pathways between both groups (Enrichment score). Pathways with Enrichment Score higher than 1.5 are shown.

In this study, a total of 992 known biochemicals were detected in the blood plasma samples. Because significant BMI differences were detected between aniridia and control groups, ANCOVA co-variate analysis was performed to assess the statistical significance of altered metabolites considering the two variates: “group” (control or aniridia patient) and “BMI” (Body max index). Through this analysis, 99 compounds were found significantly altered (p ≤ 0.05) between groups, with 53 compounds increased and 46 decreased in aniridia vs control samples. A further 75 compounds were found trending to significance (0.05 < p < 0.10), of which 39 were increased and 36 decreased in aniridia samples compared to controls. The distribution of differential metabolite classes is shown in Figure 1C, with lipids (53%) being the predominant class, followed by amino acids (23%) and nucleotides (10%).

Pathway enrichment analysis revealed significant perturbation of multiple metabolic networks between aniridia patients and healthy controls, particularly in phospholipid metabolism, purine metabolism, alanine and aspartate metabolism and nicotinamide metabolism (Figure 1D).

#### 2.1. (Phospho) Lipid metabolism disruption in aniridia patients

Glycerophospholipids are significantly reduced in the plasma from aniridia patients, particularly glycerophosphoethanolamine (GPE), glycerophosphorylcholine (GPC) (Fold change [FC] 0.80, *p* ≤ 0.01), and glycerophosphoserine (GPS) (Figure 2A). This effect is even more evident in lysophospholipids (Lyso-PL) levels, where most Lyso-PLs containing 16- and 18-carbon chain ethanolamine-, serine-, phosphatidic acid-, and even choline-species were detected in significant lower levels in aniridia patients (Figure 2A).

**Figure 2.**
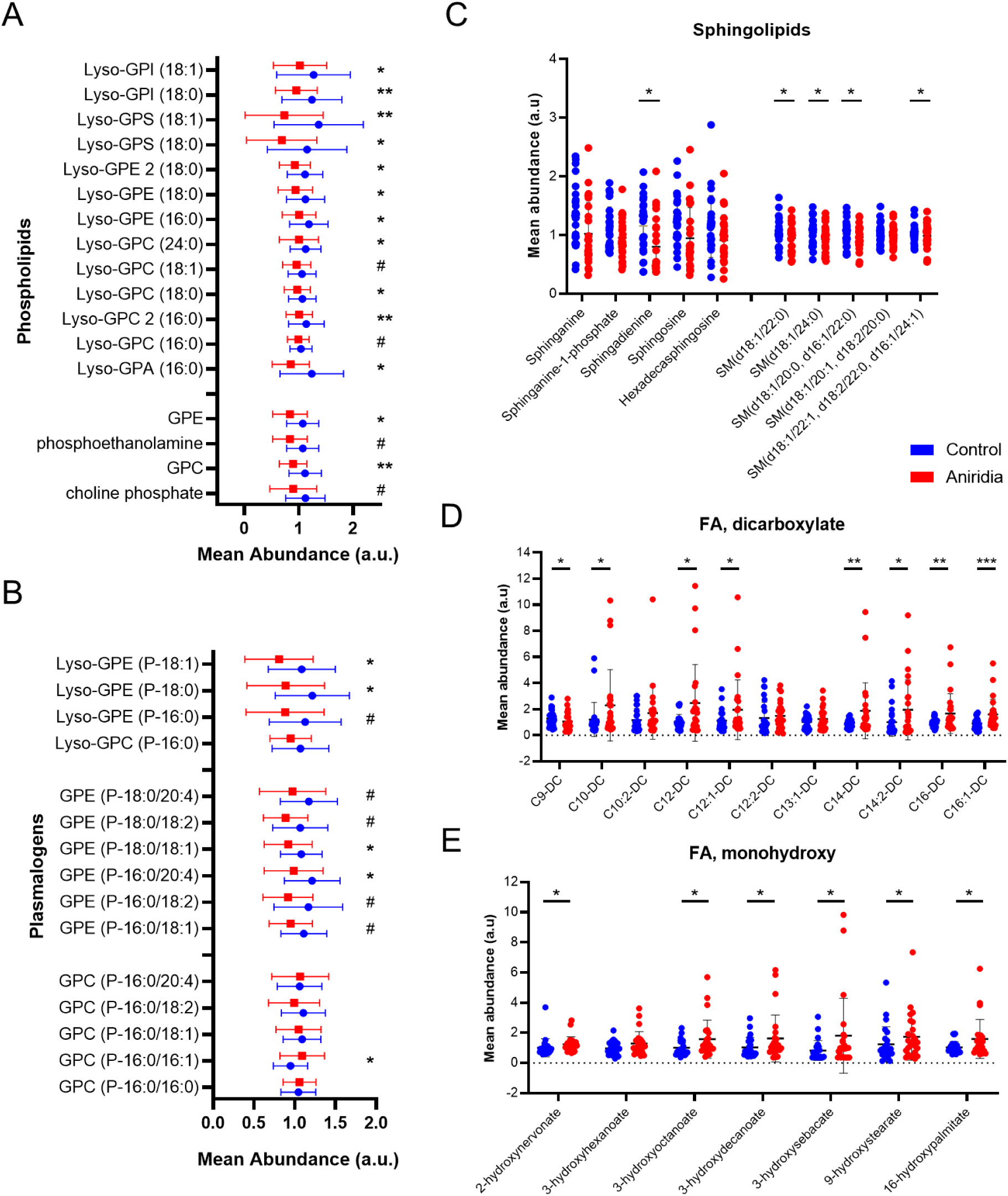
Lipid metabolism changes in aniridia patients plasma samples. A) Different phospholipid and lyso-phospholipid classes detected in control (blue) and aniridia (red) groups. B) Plasmalogens and lyso-plasmalogens metabolites in control (blue) versus aniridia (red). C) Key sphingolipids differentially detected between control (blue) and aniridia (red) groups. D, E) Levels of dicarboxylated and monohydroxylated fatty acids (FA) in both groups, respectively. n=25 per group; * p ≤ 0.05; **, p ≤ 0.01; ^#^ 0.05< p < 0.1. Abbreviations: FA, fatty acids; a.u., arbitrary units

Plasmalogens (Pls) represent a unique class of phospholipids as they contain a fatty alcohol with a vinyl-ether bond and are enriched in polyunsaturated fatty acids. The two major kinds of plasmalogens are choline Pls (PlsCho), more prevalent in heart tissue, and ethanolamine Pls (PlsEtn), mostly concentrated in myelin, hence highly abundant in the adult CNS [40]. In our study, both PlsEtn and Lyso-PlsEtn are nearly all decreased among aniridia patients compared to age matched controls (Figure 2B). In contrast, no changes were detected in level of PlsCho or Lyso-PlsCho, the exception being elevated levels of GPC (P-16:0/16:1) (FC 1.15, *p* < 0.05) (Figure 2B).

Several shingolipids were also significantly lowered in aniridia patients plasma i.e. sphingadienine (FC 0.66, p < 0.05), several 18 carbon-chain sphingomyelins (SM), like SM(d18:1/22:0) or SM(d18:1/24:0), as well as trend reduction in sphinganine (FC 0.77, *p* < 0.1), sphingosine (FC 0.78, p<0.1) and sphinganine-1-phosphate (FC 0.84, p < 0.1 (Figure 2C).

#### 2.2. Fatty acid (FA) oxidation is disrupted in aniridia patients

Aniridia patients also show significant alterations in fatty acid (FA) metabolism, particularly in FA oxidation. Dicarboxylic acids are produced via omega-oxidation in the peroxisomes and have been shown to accumulate when there are disruptions in mitochondrial beta-oxidation. Accordingly, several (medium and long chain) dicarboxylic FAs were largely increased in aniridia patients (Figure 2D). Also hydroxylated FAs, mainly monohydroxy, were significantly elevated in these patients (Figure 2E). Differences in phospholipids and carboxylated fatty acids seem to be more prevalent in those with C14, C16 and C18 species-myristate, palmitate and stearate, whilst longer chain FAs (C>18) were mostly unchanged between both groups (Supplementary Figure 2).

Increased levels of carnitines were also detected, particularly acetylcarnitine (C2) (FC 1.23, p<0.05), phenylacetylcarnitine (FC 1.72, p<0.05) and 3-decenoylcarnitine (FC 1.41, p<0.05) (Supplementary Figure 3), proving widespread lipid metabolism disruption in aniridia patients plasma.

#### 2.3. Aniridia patients have increased ketone bodies and energy changes without glucose levels alteration

High levels of dicarboxylic and 3-hydroxylated FAs, coupled with increased presence of ketone bodies are markers for ketoacidosis. Among the aniridia group, ketone bodies acetoacetate (FC 2.18, *p* < 0.05) and 3-hydroxybutyrate (BHBA) (FC 2.68, p<0.05) were elevated (Figure 3A,B). Ketone body production occurs in response to decreased carbohydrates, particularly glucose (or glycogen), or increased FAs levels in the blood; however, both glucose and 1,5- anhydroglucitol (1,5-AG) plasma levels, the latter a marker for short-term glycaemic control, were not significantly altered between both groups (Figure 3C,D).

**Figure 3.**
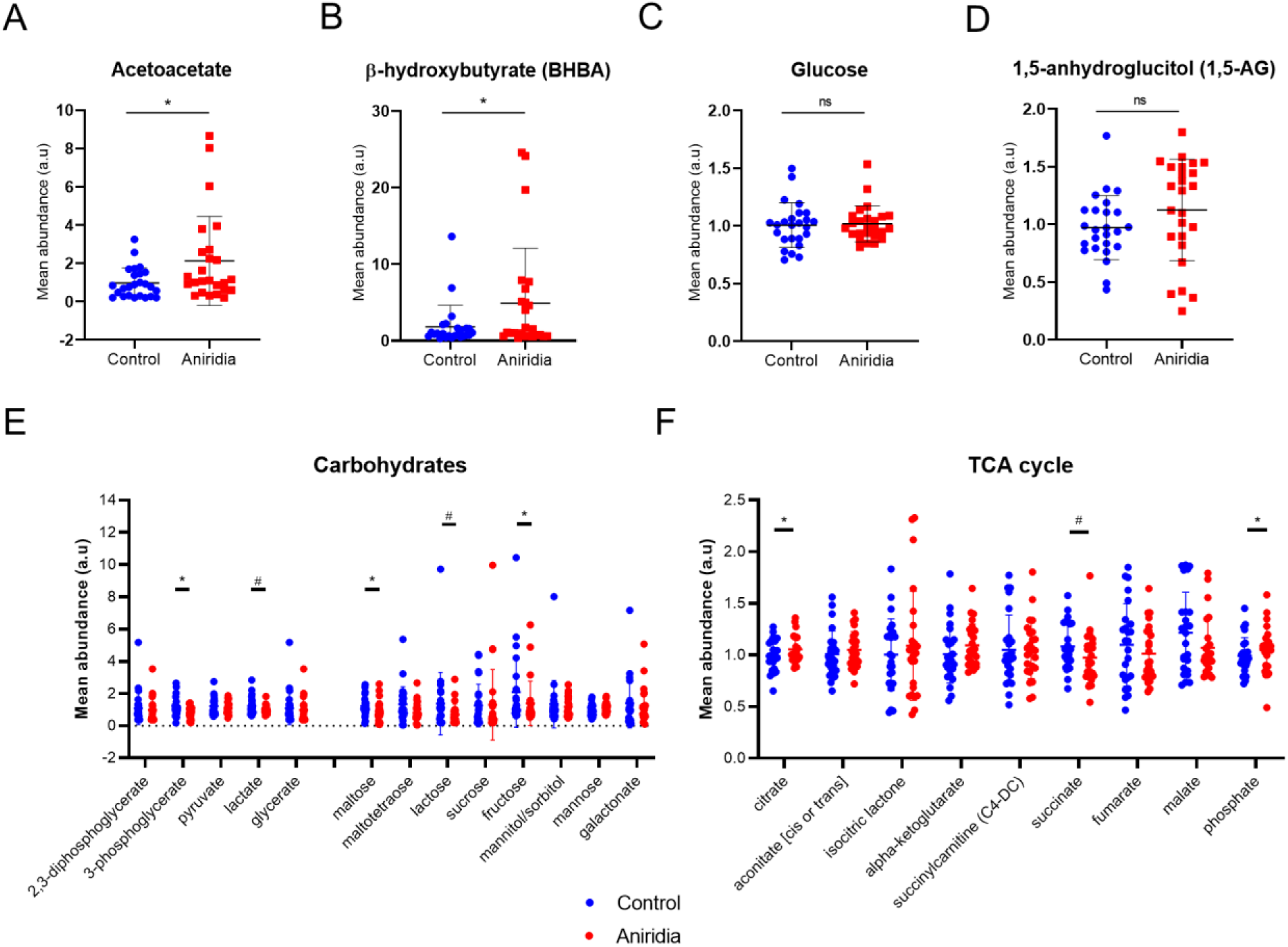
Energy metabolism alterations in aniridia patients. A-D) Box plots of Ketone bodies acetatoacetate and β-hydroxybutyrate (BHBA) as well as of glucose and glycemia marker 1,5- anhydroglucitol (1,5-AG) levels in control (blue) and aniridia (red) groups. E) Carbohydrate metabolism markers in both groups. F) TCA metabolites. n=25, * *p* ≤ 0.05; ^#^ 0.05< *p* < 0.1, ns, non-significant.

Further analysis into glycolysis, gluconeogenesis and pyruvate metabolism showed significantly decreased levels of 3-phosphoglycerate (FC 0.69, p< 0.05) as well as a trend reduction in lactate (FC 0.82, p=0.051) (Figure 3E). In addition, several carbohydrates showed reduction in aniridia patients plasma compared to controls, namely in glycogen metabolism, i.e., maltose (FC 0.61, *p*<0.05), as well as mono and polysaccharides fructose (FC 0.56, p<0.05), lactose (FC 0.58, 0.05<*p*<0.1) and sucrose (FC 0.56, 0.05<*p*<0.1) (Figure 3E). Importantly, dietary analysis showed no significant differences in intake of carbohydrates and sugars intake between both groups (Supplementary Table 2). TCA cycle metabolite analysis showed a mild but significant increase in citrate levels (FC 1.09, p<0.05) and a trend decrease in succinate (FC 0.89, 0.05<p<0.1) (Fig 3F). Pyruvate levels were not significantly altered (FC 0.90, p=0.56), unlike phosphate, which was significantly elevated (FC 1.11, p<0.05) (Figure 3F).

Results strongly support the hypothesis of mitochondrial FA oxidation defects may be occurring in aniridia. However, it must be noted that these results can be affected by differences in the fasting/fed state of the subjects at the time of blood draw.

#### 2.4. Indicators of increased oxidative stress in aniridia patients

Aniridia patients showed several altered metabolites within the methionine/ cysteine/ taurine metabolism pathways, which indicate increased demand for glutathione (Figure 4A). We found significantly elevated levels of cystathionine product alpha-ketobutyrate (FC 1.37, *p*=0.05) (Fig 4B), coupled with a trend increase in cysteine (FC 1.13, 0.05<*p*< 0.1) and a significant accumulation of by-products cystine (FC 1.11, *p*< 0.05) and cysteine-S-sulphate (FC 1.55, *p*< 0.05) (Fig 4B). In contrast, glutathione-derivatives cysteine-glutathione disulfide (FC 0.65, *p* < 0.05) and γ-glutamylalanine (FC 0.74, p<0.05) were significantly reduced in these patients, with several more γ-glutamylaminoacids trending to reduction (Figure 4B).

**Figure 4.**
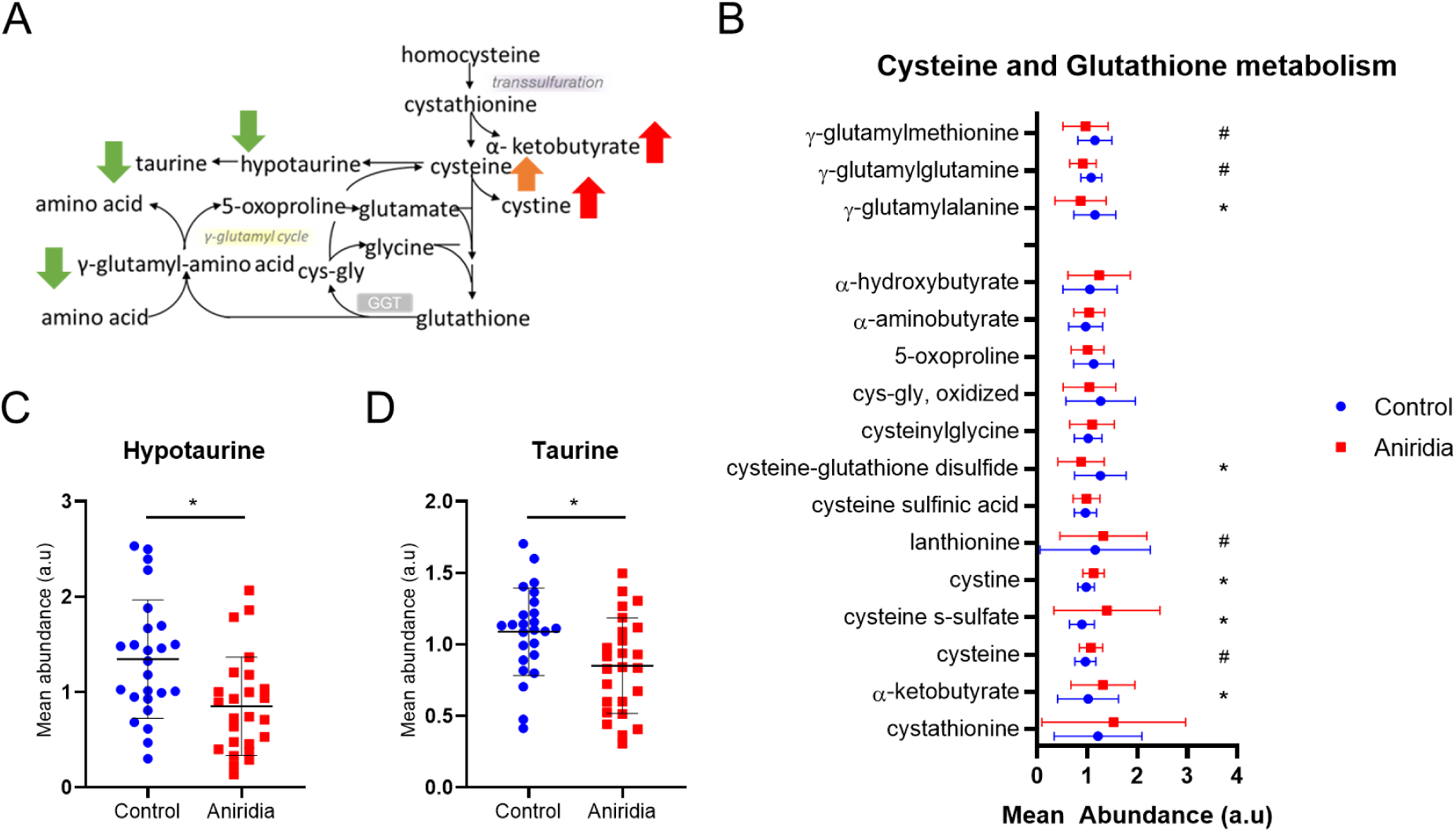
Cysteine and glutathione metabolism. A) Schematic with cysteine and glutathione pathways; arrow indicate altered expression of different metabolites in aniridia samples: significantly reduced (green), significantly elevated (red) and trend increase (orange). B) Individual metabolites from the same pathway in control (blue) and aniridia (red) samples. C,D) Box plots of metabolites hypotaurine and taurine. n=25, * p ≤ 0.05; ** p ≤ 0.01; ^#^ 0.05< p < 0.1.

In parallel, neurotransmitter and antioxidant amino acid taurine (FC 0.76, *p* < 0.05) and its precursor hypotaurine (FC 0.68, *p* < 0.05) were significantly reduced in aniridia patients (Figure 4C,D), suggesting possible shift in cysteine metabolism due to increased glutathione demand in aniridia patients, likely caused by increased oxidative stress. Glutathione levels were not detected in our study.

#### 2.5. Possible links to gut microbiome changes

##### 2.5.1. Tryptophan metabolism

Aniridia patients also displayed altered levels of tryptophan-derived metabolites. Serotonin is a tryptophan-derived hormone that has roles in mediating sleep, mood, gut movement, growth, and learning. Although a trend reduction was observed, serotonin changes were not significant in aniridia patients following ANCOVA analysis with BMI consideration (FC 0.62, p =0.18) (Figure 5A). Indoles represent a wide group of gut bacteria-derived compounds produced from tryptophan. We found that indolelactate was slightly reduced (FC 0.85, p<0.1), whilst indole-3-carboxylate was significantly increased in the aniridia cohort (FC 1.55, p<0.01) (Figure 5A).

**Figure 5.**
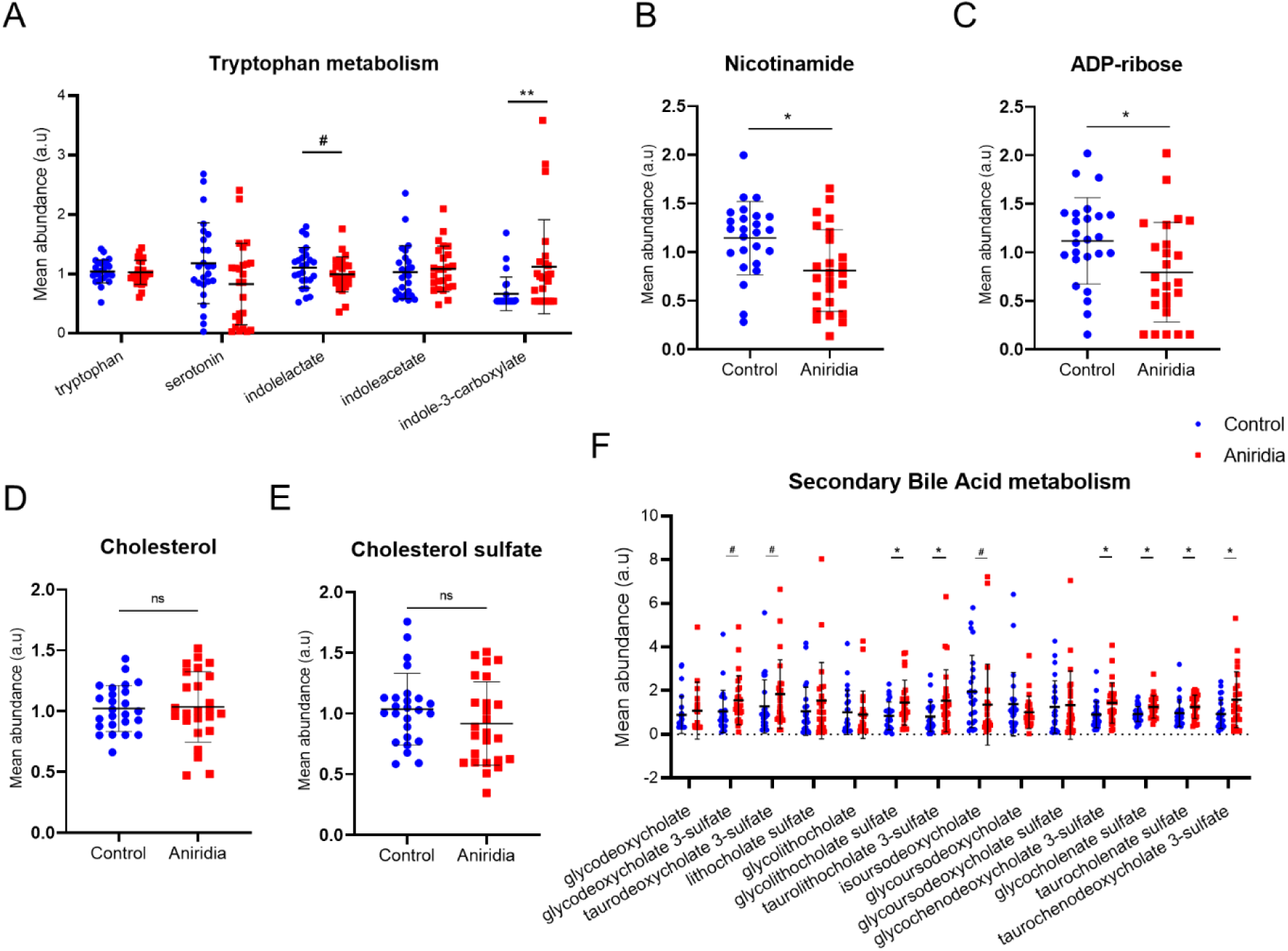
Changes in metabolites linked to gut microbiome. A) Metabolites from tryptophan metabolism pathway showing altered indole levels in aniridia (red) versus control (blue) groups. B, C) Significantly altered metabolites in the nicotinamide metabolism pathway (downstream in the tryptophan metabolism pathway). D,E) Cholesterol and cholesterol sulphate levels were unchanged between control (blue) and aniridia (red). F) Secondary bile acid levels in control (blue) and aniridia (red) groups. n=25, * p<0.05; ** p<0.01; ^#^ 0.05<p<0.1; ^ns^ not significant. Black lines represent mean and standard deviations.

Tryptophan metabolism can be diverted from serotonin production to the kynurenine pathway, producing quinolinate, which in turn is necessary for nicotinamide production. Metabolites along the kynurenine axis were unaltered but nicotinamide levels were significantly reduced in aniridia patients (FC 0.69, p<0.05) as well as adenosine 5’-diphosphoribose (ADP-ribose), an intermediate in NAD metabolism (FC 0.61, p<0.05) (Figure 5B,C). These results support the previous evidence that redox metabolism is affected in aniridia.

##### 2.5.2. Bile acid metabolism

Primary bile acids (BAs) are cholesterol catabolites mainly synthesised in the liver. Following synthesis, they are conjugated to either taurine or glycine, forming bile salts and secreted into the bile. Secondary BAs are generated when gut microbiota modify primary BAs in the colon [41]. Cholesterol and cholesterol-sulphate levels were similar between aniridia and control groups (Figure 5D,E), although small perturbations in steroid metabolism were detected in aniridia patients, such as increased levels in 3β-hydroxy-5-cholenoic acid (FC 1.42, p<0.01) and trend increase in 7-Hoca (FC 1.17, p=0.055) (Supplementary Figure 4A). Furthermore, whilst primary BA levels seemed unaltered (Supplementary Figure 4B), striking changes were detected in the levels of secondary BAs, particularly their sulphated forms: tauro- and glycodeoxycholate 3-sulfate, tauro- and glycolithocholate 3-sulfate and tauro- and glycocholenate sulfate, were all significantly increased in aniridia patients plasma (Figure 5F). Although plasma taurine was significantly reduced in aniridia (Figure 4D), no major differences between levels of taurine- or glycine-conjugated BAs were detected.

We found several other altered metabolites linked to gut microbiome that were altered in the aniridia group: trend increase in p-cresol sulphate (a microbial metabolite linked to cardiovascular disease and oxidative stress, FC 1.58, 0.05<p<0.1), and significant increased levels of 4-Hydroxychlorothalonil (FC 1.49, p<0.05) and 2’-O-methyluridine (FC 1.24, p<0.05), two key metabolites identified as predictors of gut microbiome diversity and health [42, 43] (Supplementary Figure 4C).

#### 2.6. Neuroactive metabolites

Alanine and Aspartate metabolism pathway was significantly altered in aniridia patients (Figure 6A). Alanine levels were unchanged but the derivative N,N-dimethylalanine is significantly elevated (FC 1.38, p<0.05) (Figure 6B). Importantly, aspartate levels are decreased in the aniridia group (FC 0.81, p<0.05), coupled with accumulation of downstream metabolite N-acetylaspartate (NAA) (FC 1.17, p<0.01) (Figure 6C,D). NAA is the precursor of N-acetylaspartylglutamate (NAAG), one of the most prevalent neurotransmitters in humans; however, NAAG levels were not significantly changed (FC 0.95, p=0.5)(Figure 6E). Asparagine levels, another product of aspartate, were not significant after BMI adjustment (FC 0.93, p=0.128) but the amount of its derivatives N-acetylasparagine (FC 1.33, p< 0.1) and hydroxyasparagine (FC 1.12, p<0.05) were also increased (Figure 6F-H). Lastly, β- citrylglutamate, a metabolite produced by the same enzyme capable of producing NAAG, showed a trend reduction in aniridia patients (FC 0.77, p < 0.1) (Figure 6I).

**Figure 6.**
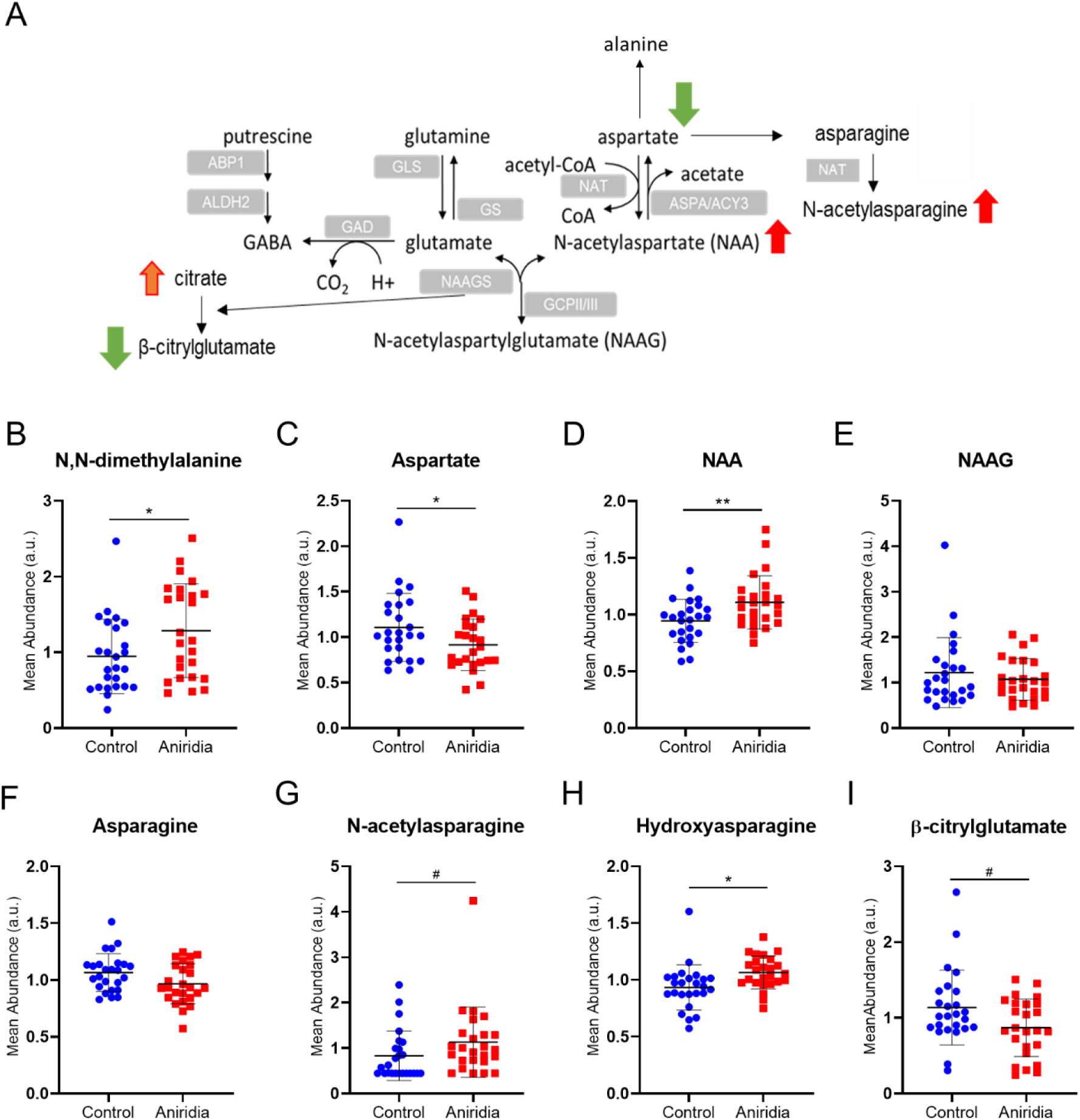
Alanine and Aspartate metabolism; A) Scheme of aspartate metabolism and neurotransmitters N-acetylaspartylglutamate (NAAG) and gamma-aminobutyric acid (GABA) production. B-I) Box plots of altered metabolites in this pathway. n=25, * p ≤ 0.05; ** p ≤ 0.01; ^#^ 0.05<p<0.1. NAA, N-acetylaspartate; NAAG, N-acetylaspartylglutamate.

Lastly, metabolites from both purine (Figure 7A-C) and pyrimidine (Figure 7D-F) metabolisms were also significantly altered in AN patients plasma, particularly inosine 5’-monophosphate (IMP, FC 0.58 p<0.05), adenosine 5’-monophosphate (AMP, FC 0.63, p<0.01), and orotate (FC 0.80, p<0.05) were significantly reduced, whilst many secondary derivatives were increased. These pathways are needed for DNA and RNA synthesis and energy production and have been implicated in neuronal development.

**Figure 7.**
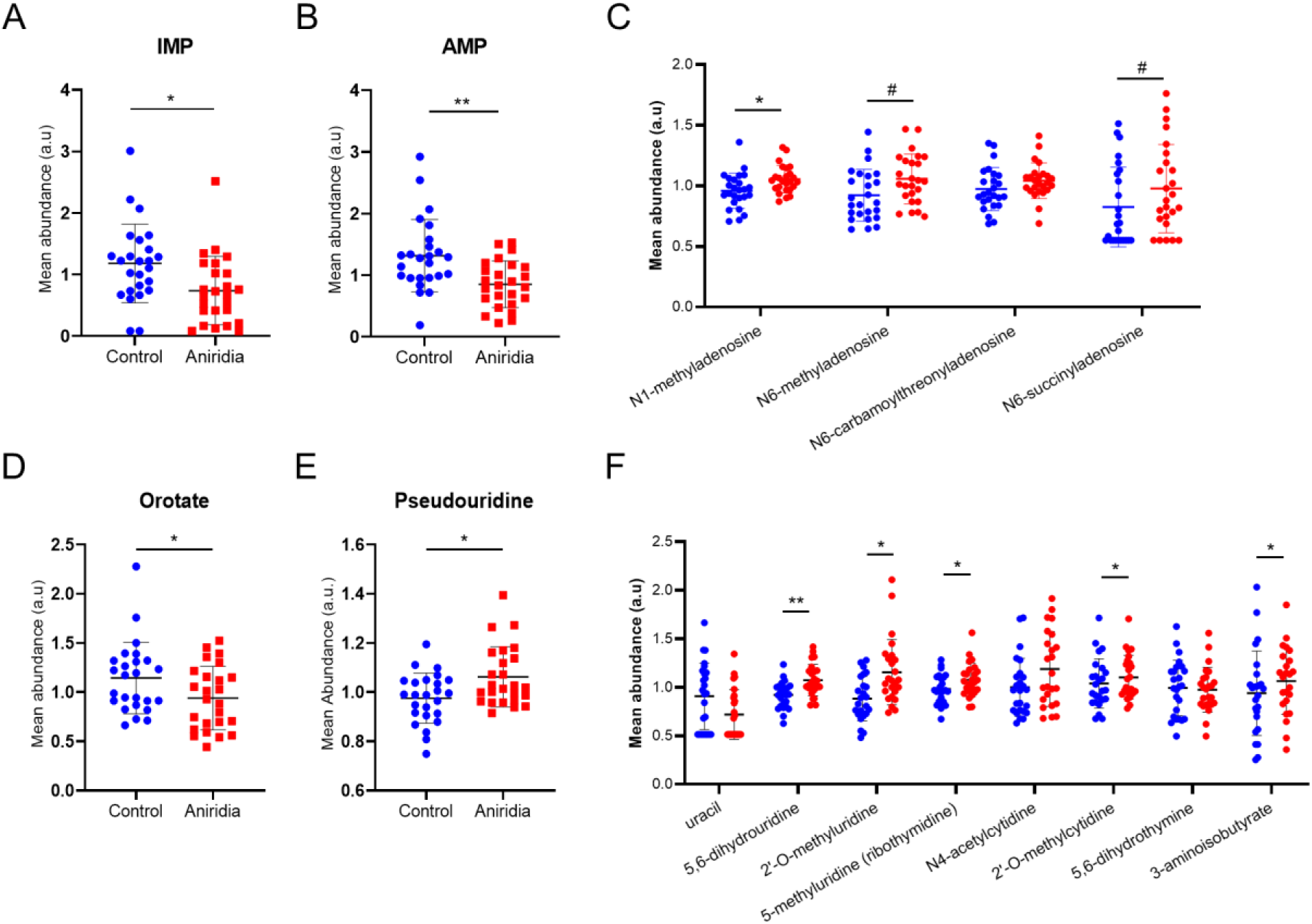
Purine and Pyrimidine metabolism. (A-C) Metabolites from purine metabolism pathway showing significantly altered levels in aniridia (red) versus control (blue) groups. (D-F) Metabolites from pyrimidine metabolism pathway showing significantly altered levels in aniridia (red) versus control (blue) groups. n=25, * p ≤ 0.05; ** p ≤ 0.01; ^#^ 0.05<p<0.1. Abbreviations: IMP, inosine 5’- monophosphate; AMP, adenosine 5’-monophosphate.

## Discussion

*PAX6* encodes a transcription factor essential for the development of the eye, pancreas and brain. Most patients with pathogenic heterozygous *PAX6* variants present with typical isolated aniridia, but a broad range of associated systemic clinical features co-exist. The effect of reduced *PAX6* levels on the metabolic function of aniridia patients has been poorly characterised. This study presents the first comprehensive metabolomic analysis of aniridia patients with molecularly confirmed *PAX6* haploinsufficiency, providing extensive evidence of plasma biomarkers associated with oxidative stress, insulin resistance and overall metabolic deregulation, as well as oxidative stress and neurotransmitter imbalance.

The metabolomic plasma profile of aniridia patients evidenced increased oxidative stress responses and redox dysregulation. In aniridia, metabolites upstream of glutathione synthesis like cystine, alpha-ketobutyrate or cysteine-S-sulphate were found in high levels and several downstream glutathione derivatives like cysteine-glutathione disulfide and gamma glutamyl amino acids were decreased. This imbalance points to a higher demand for glutathione, consistent with increased oxidative stress. Levels of taurine and its precursor hypotaurine are also reduced in our aniridia cohort; taurine is produced from cysteine and acts as an inhibitory neurotransmitter with neuroprotective and antioxidant properties crucial for brain and eye development [44, 45]. Topical taurine showed significant antioxidant effect and prevented ocular surface damage in rabbit model of dry eye disease [46]. Aside from the antioxidant properties described above, taurine is also involved in bile acid biosynthesis and can stimulate insulin release, which also seem affected in our cohort (discussed below) [44]. Altered plasma taurine levels have been associated with neurological disorders like depression, schizophrenia or autism spectrum disorder (ASD), but also with metabolic diseases like hypertension, obesity, or hypothyroidism, most of which were described in our aniridia cohort [44]. Accordingly, taurine supplementation showed promising benefits against several of these diseases, so its therapeutic potential for aniridia patients should be assessed (reviewed in [44]).

The cornea is particularly sensitive to oxidative stress due to its superficial location and exposure to external insults e.g. UV light, and deficient management of oxidative stress is one of the pathomechanisms behind several corneal disorders [47]. *Pax6* heterozygous mice corneal cells showed decreased expression of *glutathione-S-transferase* and are more susceptible to stress, which can be ameliorated by treatment with glutathione [48]. Furthermore, in the brain, Pax6 was shown to bind to promoters of *SOD1* and *CAT* genes, which code for two important reactive oxygen species (ROS) scavengers [49]. The regulatory effect of PAX6 in the expression of ROS scavengers may therefore be expanded to the extra-ocular tissues, which could explain the altered oxidative stress markers found in patients plasma.

Nicotinamide (NAM) is essential for nicotinamide adenine dinucleotide (NAD+) production and is decreased in aniridia patients, pointing to overall redox disruption. Levels of NAM-derivative NAD+ are decreased in a mouse model of glaucoma, where authors proposed it would cause mitochondrial dysfunction, resulting in increased ROS and DNA damage within retinal ganglion cells (RGCs). Furthermore, oral supplementation with NAM not only prevented but also improved RGCs survival and inner retinal function [50, 51]. Similar results were shown in humans, where NAM levels were significantly lower in the plasma of 34 primary open-angle glaucoma patients and, recently, a clinical trial with 57 glaucoma patients showed improved inner retinal function in 23% patients after oral NAM supplementation, suggesting its use as a future therapeutic strategy for glaucoma [37, 52]. Up to two thirds of aniridia patients develop glaucoma by late childhood or early adulthood [7, 10]; in our cohort, 32% (8/25) of patients were diagnosed with glaucoma. Nicotinamide supplementation could be considered in aniridia as it may delay glaucoma development/ progression. Further metabolites found in common with other metabolomics glaucoma studies include decreased spermine, increased acetylcarnitine and increased decenoylcarnitine levels [53].

Phosphoetanolamines-derived plasmalogens (PLsEth) and lysoplasmalogens were largely reduced in our aniridia cohort, also potentially contributing to the altered oxidative stress status. PLsEth are very prevalent in brain myelin and the lens, where levels tend to reduce with age [40]. In fact, congenital cataracts are a key symptom in patients with defects in plasmalogen synthesis, like Zellweger syndrome or Rhizomelic Chondrodysplasia Punctata [40]. PAX6 deficiency is associated with cataract formation, with up to 85% of aniridia patients developing early onset lens opacities [7]; in our cohort, 75% (19/25) of patients had cataracts/lens opacities. Assessing whether plasma PLsEth levels correlate with those in the cataract lens would establish the potential of it being a biomarker for cataract formation/ severity in aniridia patients.

Overall lysophospholipids (Lyso-PLs) were decreased in aniridia plasma. Choline-derived Lyso-PLs have been identified as negative predictor of Type 2 diabetes progression [54], and accordingly, several were significantly decreased in aniridia patients. Lower Lyso-PLs may also reflect altered activity of phospholipase A2, which cleaves phospholipids generating Lyso-PL and a free fatty acid [55]. Increased phospholipase A2 has been associated with inflammation, schizophrenia or autism but no link to PAX6 has been reported to date [56].

Plasma levels of several dicarboxylic and hydroxy FAs were increased in AN plasma. Dicarboxylic acids are produced via omega-oxidation in the peroxisomes and have been shown to accumulate when there are disruptions in mitochondrial beta-oxidation. FA hydroxylation is mostly performed by a family of enzymes termed cytochromes P450s during lipid oxidation. P450s are monooxygenases that also oxidize steroids and xenobiotics, and are important for hormone synthesis and breakdown. Cytochrome P4501B1 (CYP1B1) has functions in adipogenesis and fatty acid oxidation and is the only P450s member we found with known relation to PAX6, as it has been reported *Cyp1b1* proximal promoter contains an alternative binding site for Pax6 in mice adipocytes [57–59]. Interestingly, *CYP1B1* is also essential for eye development, with mutations causing primary congenital glaucoma (OMIM # 231300) and anterior segment dysgenesis/ Peters Anomaly (OMIM # 617315) [58].

Our results therefore suggest that PAX6 haploinsufficiency may impact beta and omega oxidation processes in mitochondria. Altered metabolites associated with both processes may suggest that aniridia patients have some alteration to the pathway of lipid oxidation, which, combined with redox abnormalities, may extend to overall mitochondrial function. However, omega-oxidation and some beta-oxidation also take place in peroxisomes. In fact, lower plasmalogens and altered bile acid metabolism are also found in peroxisomal disorders as well as several other altered compounds linked to peroxisomal disorders: 7-alpha-hydroxy-3- oxo-4-cholestenoate (7-Hoca), increased 3-Hydroxysebacic and 3-Hydroxydecanedioic acids. We therefore hypothesize that the links to mitochondrial dysfunction in aniridia patients can also expand to (or derive from?) the peroxisome. However, it must be mentioned that these results may be affected by differences in the fasting/fed state of the subjects at the time of blood draw.

Further metabolic changes in AN patients plasma include increased presence of ketone bodies acetoacetate and BHBA, which suggest a tendency for metabolic acidosis. Accordingly, markers of inborn error of metabolism diseases linked to aciduria, such as isovaleric aciduria (OMIM #243500) or medium chain acylCoA dehydrogenase deficiency (OMIM #201450), were found increased in the aniridia cohort, i.e 3-hydroxy FAs and methylsuccinate. Although symptoms and levels described in patients with the aforementioned metabolic diseases were more severe than observed in the AN cohort, these results suggest that aniridia patients may require monitoring for acidemia symptoms.

Ketogenesis can be upregulated by hormones such as glucagon or cortisol; however, insulin is the main hormonal regulator of this process, suggesting an altered insulin state in aniridia patients included in this study. PAX6 plays an essential role in pancreas development and in the continuous regulation of insulin and glucagon synthesis in adulthood. *Pax6* deletion in mice pancreatic β cells caused increase in glucagon and ghrelin hormone levels but decrease in insulin, resulting in hyperglycemia and severe ketosis [60].

In a recent retrospective analysis of 86 aniridia patients, we showed that 13% of patients were diagnosed with T2D, which is double the UK prevalence [10]. However, clinically diagnosed diabetics were excluded from this study. Nonetheless, several metabolites associated with T2D were significantly altered in the AN group, such as increased acylcarnitines, a trend towards increases in cysteine, accumulation of ketone bodies and increased bile acids, which suggest most patients could already be in an insulin resistance/ prediabetic stage [54, 61, 62]. Plasma long-chain saturated FAs (C14:0, C16:0 and C18:0) may induce insulin resistance and are considered risk factors for T2D and metabolic syndrome [63]. Decreased plasmalogens are also associated with increased lipid oxidation and lower Pls-Etn levels have been detected in patients with metabolic syndrome, T1D and T2D, and hyperlipidemia [40, 64].

Bile acids (BA) are important modulators of lipid and glucose metabolism, and increased levels have been found in plasma of non-alcoholic fatty liver disease (NAFLD) and in serum of T2D patients [41, 65]. For example, taurolithocholate 3-sulphate, which was over 2-fold increased in our aniridia cohort, was found to induce insulin resistance in *in vitro* rat hepatocytes [66]. It is thought that BA sulphation is a counterbalance mechanism to alleviate BA accumulation by increasing BA solubility and excretion [67]. Increased BA sulphation is present in cholestasis, characterised by decreased bile flow, resulting in BA accumulation, increased ROS and hepatocytotoxicity [41]. However, the clinical consequences of high BA levels in aniridia are unclear but these results seem to point out the need for patients to be monitored for regular liver function tests.

Gut microbiome alterations in aniridia patients should also be considered. Secondary BA are produced by modification of primary BA by the gut microbiota and were particularly elevated in AN plasma [41]. Furthermore, indoles are gut microbiota-derived tryptophan metabolites and are suggested to exert significant biological effects and contribute to the aetiology of cardiovascular, metabolic, and psychiatric diseases [68]. A trend towards a decrease of indole lactate and increased indole-3-carboxylate was identified in AN plasma; indole-3-lactate has known anti-inflammatory properties and Indole-3-carboxylate was detected in uremic patients and has been found elevated in patients with liver diseases [69, 70]. However, the exact role of this compound is not known yet.

Serotonin is a hormone and neurotransmitter that regulates sleep, mood and behaviour also derived from tryptophan metabolism. Although its levels seem reduced in AN patients plasma, no statistical significance was achieved after ANCOVA analysis, pointing to serotonin levels being likely more associated with BMI. Accordingly, several reports show that reduced plasma serotonin is related to obesity, insulin resistance, diabetes and metabolic syndrome [71–75]. Other metabolites altered in aniridia patients plasma but associated to obesity/high BMI via ANCOVA analysis included decreased asparagine, increased carnitine and decreased uracil [76, 77]. PAX6 role in endocrine pancreas function and in regulation of energy metabolism has been the accepted reason behind the increased BMI detected in many aniridia patients. However, other players and mechanisms could be considered. WAGRO is a form of syndromic aniridia caused by contiguous deletion of *PAX6, WT1* and *BDNF* genes and is characterised by aniridia, Wilms tumour, genitourinary anomalies and mental retardation as well as obesity [4]. Variants affecting *BDNF* are associated with obesity and hyperphagia, with reduced BDNF levels correlating with higher BMI [5, 6]. Pax6 was shown to bind to a *Bdnf* promoter in the mouse brain and Bdnf plasma levels appeared to correlate with brain levels [49, 78]. The increased BMI in our aniridia group could therefore also be (at least partially) explained by a more direct PAX6-BDNF relationship, although the underlying mechanisms are not yet elucidated. WAGRO patients were not included in this study, but it would be interesting to measure BDNF levels in blood from isolated aniridia patients to understand if *PAX6* haploinsufficiency results in direct reduction of BDNF levels.

Several neuroactive metabolites were also significantly changed in aniridia plasma samples compared to the controls. Aspartate is a major excitatory neurotransmitter and can be decreased in depression [79]. N-acetylaspartate (NAA) is one of the most prevalent neurotransmitters in humans and is synthesized from aspartate and acetyl-coenzyme A in neuronal mitochondria, being important for myelin lipid synthesis. Elevated NAA levels are markers for Canavan disease (OMIM #271900), a neurodegenerative disease caused by autosomal recessive mutations in *ASPA* [80, 81]. There is no known relation between PAX6 and NAA but *ASPA* as well as *ASNS,* which produces asparagine from aspartate, are known targets of PAX6 [82] and patients can show vision impairment caused by optic atrophy (although optic nerve hypoplasia and retinal degeneration similar to RP have been described) [83, 84]. NAA downstream metabolite N-acetylaspartylglutamate (NAAG) seems unchanged in aniridia patients. However, levels of β-citrylglutamate, a structural analog of NAAG produced from citrate by the same enzyme, NAAGS, seem reduced. We hypothesize there could be a diversion of NAAGS activity to maintain NAAG levels at the expense of β-citrylglutamate. The biological relevance of β-citrylglutamate is still largely unexplored, but it is found in high levels in the developing brain, testes and eye, where it seems required for differentiation of lens epithelial cells [85–87]. This compound could therefore have a role in the development of cataracts in aniridia patients.

Numerous proteins and enzymes in the pathways and processes described in this study, i.e. lipid oxidation, plasmalogen synthesis, bile acid and redox metabolism, are regulated by a superfamily of ligand-activated transcription factors called Peroxisome proliferator-activated receptors (PPARs). Based on the current evidence, it could be suggested that the metabolic defects identified in AN patients could be caused not only by the PAX6 defective role in pancreas and insulin production directly but also through PPAR (or even mutual) deregulation. PPAR gamma (PPARγ) was shown to interact with PAX6 in the regulation of glucagon gene [88, 89]. Interestingly, altered *PPARγ* expression also induced an ocular phenotype in octopus, where both overexpression and knockdown inhibited *pax6* expression causing small eye phenotype and lens opacities (microencephaly and enlarged trunk was also described) [90].

Also, expression of *PPARγ* was shown to be downregulated in aniridia patients’ and *PAX6*- edited limbal stem cells [91, 92]. Although further elucidation between PAX6 and PPARs is required, It would be interesting to test the potential of PPAR agonists, particularly PPARγ, in improving the systemic metabolic (and perhaps also ocular) alterations seen in aniridia patients.

Another interesting therapeutic approach could be amlexanox, We have recently shown that this compound can increase PAX6 levels in aniridia patient-derived retinal and corneal cell models through its nonsense suppression properties [93]. Notably, amlexanox has also been shown to improve dyslipidemia and target bile acid synthesis [94] and its use ameliorated insulin resistance and obesity phenotypes in both mice and humans [95, 96].

To conclude, this study provides novel insights into the systemic derangements behind *PAX6* haploinsufficiency, and proposes unexplored molecular mechanisms that can ultimately improve clinical care of these patients. Further molecular studies are needed to understand how the metabolites and pathways described in this study are related to *PAX6* haploinsufficiency and how they can be translated into therapeutics to prevent or slow down disease progression.

### Study limitations

Untargeted metabolomic approach of aniridia patients plasma identified several potential biomarkers for common PAX6-associated systemic anomalies like obesity, diabetes, or CNS abnormalities. Nonetheless, we identified several limitations to this study that may have affected the group separation. Firstly, whilst ocular findings are clinically acquired through examination, extra-ocular findings in our patient and control cohort are dependent on self-reporting. This may result in an under-reporting of systemic/extra-ocular conditions which are either unreported by the patient or not diagnosed as the patient has not sought medical investigations; for example, hypertension, depression, diabetes and pre-diabetes are often symptomless and unless actively investigated may not be known to the patient. Next, the wide span of ages within the study, which we tried to solve by including age-matched controls. Also, blood samples were collected from non-fasting individuals, the vast majority in the mornings but in a few cases, due to proband availability, samples were collected in the afternoon. A significant challenge of this study comes from the ocular and systemic phenotypic heterogeneity present within the aniridia patients, which is naturally present in *PAX6*-related disease. We could not control the effects of medication that both patients and controls were on, and understanding how these may somehow mask or contribute to some metabolomic signals would be helpful.

## Methods

### ETHICS, CONSENT AND PERMISSION

Patients with a confirmed molecular diagnosis of PAX6-aniridia were identified from the Moorfields Eye Hospital Inherited Eye Disease Database (Moorfields Eye Hospital NHS Foundation Trust, London, UK). All study participants gave informed consent to participate, and all procedures were conducted in adherence to the tenets of the Declaration of Helsinki. The study received ethical approval from the relevant local bodies (Research Ethics Number: 12/LO/0141.

### CLINICAL EVALUATION

Electronic and paper medical records from Moorfields Eye Hospital NHS Foundation Trust, London, UK were reviewed for all patients. Systemic clinical information including height, weight, current medications and past medical history were self-reported by participants at the time of study recruitment. Ocular and self-reported systemic conditions are summarised in Supplementary Table 1.

### DIETARY INTAKE

Subjects were asked to complete a food frequency questionnaire (FFQ) on their average consumption of various foods and drinks over the past 12 months. The validated FFQ comprised a list of 147 food items where participants were asked to indicate their usual consumption from one of nine frequency categories ranging from “never or less than once per month” to “six or more times per day.” [39].

### SAMPLE COLLECTION

The study included blood plasma samples from non-fasting 25 human Aniridia patient and 25 age and gender-matched healthy individuals. Plasma was extracted as previously reported [35]. Only plasma samples that had not previously been thawed were included for metabolomics analysis. When all samples were collected, they were sent in dry ice to Metabolon Inc (Durham, NC, USA).

### METABOLOMICS ANALYSIS

Blood plasma metabolite extractions for Ultrahigh Performance Liquid Chromatography-Tandem Mass Spectroscopy (UPLC-MS/MS), quantification and preliminary biological interpretation were performed by Metabolon Inc (Durham, NC, USA), according to the protocol described in Supplementary Materials and Methods.

### STATISTICAL ANALYSIS

Differences in Age, BMI and dietary intake between both groups were calculated with Mann Whitney tests (GraphPad Inc). Metabolite profiles in Aniridia patients and healthy controls were quantified in terms of relative abundance and median scaled to 1. Following log transformation and imputation of missing values, if any, with the minimum observed value for each compound imputed, statistical analyses were performed to identify significant differences between experimental groups using ArrayStudio (Omicsoft, Cary, NC, USA), R version 2.14.2, and/or SAS v9.4 (https://www.r-project.org/). Metabolite profile distinctions between Aniridia patients and healthy individuals were initially evaluated by Welch’s two-sample t-test. Fold change (FC) was determined by dividing the relative abundance of the metabolite in the aniridia patients blood plasma by the relative abundance of the metabolite in the blood plasma of healthy control individuals. FC values with p ≤ 0.05 were considered statistically significant in this study. FC with 0.05 < p < 0.10 were considered as trending towards significance. Analysis of covariance (ANCOVA) was used for variables “Group” (patient or control) and “BMI” (body mass index). Metabolites with FC values with p ≤ 0.05 associated with “Group” variable were considered significantly altered in the Aniridia group.

## Acknowledgements and funding

We would like to thank all patients and volunteers for their participation in this study. Also Metabolon Inc., particularly Kimberley Jackson, for technical and analytical support, Cecile Mejecase (The Francis Crick Institute, London, UK) for assisting with samples preparation, and Sadaf Farooqi (University of Cambridge Metabolic Research Laboratories, Institute of Metabolic Science and NIHR Cambridge Biomedical Research Centre, Cambridge, UK) for fruitful discussions during manuscript preparation. This research was funded by the Wellcome Trust (205174/Z/16/Z) and Moorfields Eye Charity, and supported by the National Institute for Health and Care Research (NIHR) Biomedical Research Centre at Moorfields Eye Hospital NHS Foundation Trust and UCL Institute of Ophthalmology. The views expressed are those of the author(s) and not necessarily those of the NHS, the NIHR or the Department of Health and Social Care.

## Author contributions

Conceptualisation: MM; Data collection: DLC, VK, JS; Data analysis – DLC, VK, JS, AAW; Visualisation: DLC; Funding acquisition: MM; Project administration: MM; Writing – first draft: DLC; Writing – review and editing – DLC, VK, MM. All authors have read and approved the submitted version

## References

1. Ton, C.C., et al., Positional cloning and characterization of a paired box- and homeobox-containing gene from the aniridia region. Cell, 1991. 67(6): p. 1059–74.

2. Moosajee, M., M. Hingorani, and A.T. Moore, PAX6-Related Aniridia, in GeneReviews(®). 2018, University of Washington, Seattle: Seattle (WA).

3. Riccardi, V.M., et al., Chromosomal imbalance in the Aniridia-Wilms’ tumor association: 11p interstitial deletion. Pediatrics, 1978. 61(4): p. 604–10.

4. Han, J.C., et al., Brain-derived neurotrophic factor and obesity in the WAGR syndrome. N Engl J Med, 2008. 359(9): p. 918–27.

5. Sandrini, L., et al., Association between Obesity and Circulating Brain-Derived Neurotrophic Factor (BDNF) Levels: Systematic Review of Literature and Meta-Analysis. Int J Mol Sci, 2018. 19(8).

6. Xu, B. and X. Xie, Neurotrophic factor control of satiety and body weight. Nat Rev Neurosci, 2016. 17(5): p. 282–92.

7. Lima Cunha, D., et al., The Spectrum of PAX6 Mutations and Genotype-Phenotype Correlations in the Eye. Genes (Basel), 2019. 10(12).

8. Lagali, N., et al., PAX6 Mutational Status Determines Aniridia-Associated Keratopathy Phenotype. Ophthalmology, 2020. 127(2): p. 273–275.

9. Hingorani, M., et al., Detailed ophthalmologic evaluation of 43 individuals with PAX6 mutations. Invest Ophthalmol Vis Sci, 2009. 50(6): p. 2581–90.

10. Kit, V., et al., Longitudinal genotype-phenotype analysis in 86 PAX6-related aniridia patients. JCI Insight, 2021.

11. Berntsson, S.G., et al., Aniridia with PAX6 mutations and narcolepsy. J Sleep Res, 2020. 29(6): p. e12982.

12. Sisodiya, S.M., et al., PAX6 haploinsufficiency causes cerebral malformation and olfactory dysfunction in humans. Nat Genet, 2001. 28(3): p. 214–6.

13. Yasuda, T., et al., PAX6 mutation as a genetic factor common to aniridia and glucose intolerance. Diabetes, 2002. 51(1): p. 224–30.

14. Boese, E.A., et al., Novel Intragenic PAX6 Deletion in a Pedigree with Aniridia, Morbid Obesity, and Diabetes. Curr Eye Res, 2020. 45(1): p. 91–96.

15. Motoda, S., et al., Case of a novel PAX6 mutation with aniridia and insulin-dependent diabetes mellitus. J Diabetes Investig, 2019. 10(2): p. 552–553.

16. Tian, W., et al., Heterozygous PAX6 mutations may lead to hyper-proinsulinaemia and glucose intolerance: A case-control study in families with congenital aniridia. Diabet Med, 2021. 38(2): p. e14456.

17. Panneerselvam, A., et al., PAX proteins and their role in pancreas. Diabetes Res Clin Pract, 2019. 155: p. 107792.

18. Ashery-Padan, R., et al., Conditional inactivation of Pax6 in the pancreas causes early onset of diabetes. Dev Biol, 2004. 269(2): p. 479–88.

19. Sander, M., et al., Genetic analysis reveals that PAX6 is required for normal transcription of pancreatic hormone genes and islet development. Genes Dev, 1997. 11(13): p. 1662–73.

20. St-Onge, L., et al., Pax6 is required for differentiation of glucagon-producing alpha-cells in mouse pancreas. Nature, 1997. 387(6631): p. 406–9.

21. Hart, A.W., et al., The developmental regulator Pax6 is essential for maintenance of islet cell function in the adult mouse pancreas. PLoS One, 2013. 8(1): p. e54173.

22. Singer, R.A., et al., The Long Noncoding RNA Paupar Modulates PAX6 Regulatory Activities to Promote Alpha Cell Development and Function. Cell Metab, 2019. 30(6): p. 1091–1106.e8.

23. Mitchell, R.K., et al., The transcription factor Pax6 is required for pancreatic β cell identity, glucose-regulated ATP synthesis, and Ca(2+) dynamics in adult mice. J Biol Chem, 2017. 292(21): p. 8892–8906.

24. Netland, P.A., et al., Ocular and systemic findings in a survey of aniridia subjects. J aapos, 2011. 15(6): p. 562–6.

25. Macdonald, G.C., et al., Deletion distal to the PAX6 coding region reveals a novel basis for familial cosegregation of aniridia and diabetes mellitus. Diabetes Res Clin Pract, 2019. 148: p. 64–71.

26. Nishi, M., et al., A case of novel de novo paired box gene 6 (PAX6) mutation with early-onset diabetes mellitus and aniridia. Diabet Med, 2005. 22(5): p. 641–4.

27. Wen, J.H., et al., Paired box 6 (PAX6) regulates glucose metabolism via proinsulin processing mediated by prohormone convertase 1/3 (PC1/3). Diabetologia, 2009. 52(3): p. 504–13.

28. Duan, D., et al., Spatiotemporal expression patterns of Pax6 in the brain of embryonic, newborn, and adult mice. Brain Struct Funct, 2013. 218(2): p. 353–72.

29. Grant, M.K., et al., Structural and functional consequences of PAX6 mutations in the brain: Implications for aniridia. Brain Res, 2021. 1756: p. 147283.

30. Kikkawa, T., et al., The role of Pax6 in brain development and its impact on pathogenesis of autism spectrum disorder. Brain Res, 2019. 1705: p. 95–103.

31. Bamiou, D.E., et al., Deficient auditory interhemispheric transfer in patients with PAX6 mutations. Ann Neurol, 2004. 56(4): p. 503–9.

32. Hanish, A.E., et al., Pineal hypoplasia, reduced melatonin and sleep disturbance in patients with PAX6 haploinsufficiency. J Sleep Res, 2016. 25(1): p. 16–22.

33. Bonelli, R., et al., Systemic lipid dysregulation is a risk factor for macular neurodegenerative disease. Sci Rep, 2020. 10(1): p. 12165.

34. Laíns, I., et al., Human Plasma Metabolomics in Age-Related Macular Degeneration: Meta-Analysis of Two Cohorts. Metabolites, 2019. 9(7).

35. Lima Cunha, D., et al., REP1-deficiency causes systemic dysfunction of lipid metabolism and oxidative stress in choroideremia. JCI Insight, 2021.

36. Chao de la Barca, J.M., et al., A Plasma Metabolomic Profiling of Exudative Age-Related Macular Degeneration Showing Carnosine and Mitochondrial Deficiencies. J Clin Med, 2020. 9(3).

37. Kouassi Nzoughet, J., et al., Nicotinamide Deficiency in Primary Open-Angle Glaucoma. Invest Ophthalmol Vis Sci, 2019. 60(7): p. 2509–2514.

38. Bingham, S.A., et al., Validation of dietary assessment methods in the UK arm of EPIC using weighed records, and 24-hour urinary nitrogen and potassium and serum vitamin C and carotenoids as biomarkers. Int J Epidemiol, 1997. 26 Suppl 1: p. S137–51.

39. Welch, A.A., et al., The CAFE computer program for nutritional analysis of the EPIC-Norfolk food frequency questionnaire and identification of extreme nutrient values. J Hum Nutr Diet, 2005. 18(2): p. 99–116.

40. Braverman, N.E. and A.B. Moser, Functions of plasmalogen lipids in health and disease. Biochim Biophys Acta, 2012. 1822(9): p. 1442–52.

41. Ahmad, T.R. and R.A. Haeusler, Bile acids in glucose metabolism and insulin signalling - mechanisms and research needs. Nat Rev Endocrinol, 2019. 15(12): p. 701–712.

42. Wilmanski, T., et al., Blood metabolome predicts gut microbiome α-diversity in humans. Nat Biotechnol, 2019. 37(10): p. 1217–1228.

43. Lin, C.J., et al., Serum p-cresyl sulfate predicts cardiovascular disease and mortality in elderly hemodialysis patients. Arch Med Sci, 2013. 9(4): p. 662–8.

44. Jakaria, M., et al., Taurine and its analogs in neurological disorders: Focus on therapeutic potential and molecular mechanisms. Redox Biol, 2019. 24: p. 101223.

45. Tao, Y., et al., Systemic taurine treatment provides neuroprotection against retinal photoreceptor degeneration and visual function impairments. Drug Des Devel Ther, 2019. 13: p. 2689–2702.

46. Bucolo, C., et al., Antioxidant and Osmoprotecting Activity of Taurine in Dry Eye Models. J Ocul Pharmacol Ther, 2018. 34(1-2): p. 188–194.

47. Vallabh, N.A., V. Romano, and C.E. Willoughby, Mitochondrial dysfunction and oxidative stress in corneal disease. Mitochondrion, 2017. 36: p. 103–113.

48. Ou, J., et al., Chronic wound state exacerbated by oxidative stress in Pax6+/- aniridia-related keratopathy. J Pathol, 2008. 215(4): p. 421–30.

49. Maurya, S.K. and R. Mishra, Pax6 Binds to Promoter Sequence Elements Associated with Immunological Surveillance and Energy Homeostasis in Brain of Aging Mice. Ann Neurosci, 2017. 24(1): p. 20–25.

50. Williams, P.A., et al., Vitamin B(3) modulates mitochondrial vulnerability and prevents glaucoma in aged mice. Science, 2017. 355(6326): p. 756–760.

51. Williams, P.A., et al., Nicotinamide treatment robustly protects from inherited mouse glaucoma. Commun Integr Biol, 2018. 11(1): p. e1356956.

52. Hui, F., et al., Improvement in inner retinal function in glaucoma with nicotinamide (vitamin B3) supplementation: A crossover randomized clinical trial. Clin Exp Ophthalmol, 2020. 48(7): p. 903–914.

53. Wang, Y., et al., Metabolomics in Glaucoma: A Systematic Review. Invest Ophthalmol Vis Sci, 2021. 62(6): p. 9.

54. Gall, W.E., et al., *alpha-hydroxybutyrate is an early biomarker of insulin resistance and glucose intolerance in a nondiabetic population*. PLoS One, 2010. 5(5): p. e10883.

55. Hermansson, M., K. Hokynar, and P. Somerharju, Mechanisms of glycerophospholipid homeostasis in mammalian cells. Prog Lipid Res, 2011. 50(3): p. 240–57.

56. Bell, J.G., et al., Essential fatty acids and phospholipase A2 in autistic spectrum disorders. Prostaglandins Leukot Essent Fatty Acids, 2004. 71(4): p. 201–4.

57. Larsen, M.C., et al., Cytochrome P450 1B1: An unexpected modulator of liver fatty acid homeostasis. Arch Biochem Biophys, 2015. 571: p. 21–39.

58. Maguire, M., et al., Cyp1b1 directs Srebp-mediated cholesterol and retinoid synthesis in perinatal liver; Association with retinoic acid activity during fetal development. PLoS One, 2020. 15(2): p. e0228436.

59. Zheng, W., et al., Stimulation of mouse Cyp1b1 during adipogenesis: characterization of promoter activation by the transcription factor Pax6. Arch Biochem Biophys, 2013. 532(1): p. 1–14.

60. Swisa, A., et al., PAX6 maintains β cell identity by repressing genes of alternative islet cell types. J Clin Invest, 2017. 127(1): p. 230–243.

61. de Mello, V.D., et al., Indolepropionic acid and novel lipid metabolites are associated with a lower risk of type 2 diabetes in the Finnish Diabetes Prevention Study. Sci Rep, 2017. 7: p. 46337.

62. Ferrannini, E., et al., Early metabolic markers of the development of dysglycemia and type 2 diabetes and their physiological significance. Diabetes, 2013. 62(5): p. 1730–7.

63. A, I.S.S., A.B. C, and J.S. A, Changes in Plasma Free Fatty Acids Associated with Type-2 Diabetes. Nutrients, 2019. 11(9).

64. Colas, R., et al., LDL from obese patients with the metabolic syndrome show increased lipid peroxidation and activate platelets. Diabetologia, 2011. 54(11): p. 2931–40.

65. Haeusler, R.A., et al., Human insulin resistance is associated with increased plasma levels of 12α-hydroxylated bile acids. Diabetes, 2013. 62(12): p. 4184–91.

66. Mannack, G., et al., Taurolithocholic acid-3 sulfate impairs insulin signaling in cultured rat hepatocytes and perfused rat liver. Cell Physiol Biochem, 2008. 21(1-3): p. 137–50.

67. Alnouti, Y., Bile Acid sulfation: a pathway of bile acid elimination and detoxification. Toxicol Sci, 2009. 108(2): p. 225–46.

68. Su, X., Y. Gao, and R. Yang, Gut Microbiota-Derived Tryptophan Metabolites Maintain Gut and Systemic Homeostasis. Cells, 2022. 11(15).

69. Byrd, D.J., et al., Indolic tryptophan metabolism in uraemia. Proc Eur Dial Transplant Assoc, 1976. 12: p. 347–54.

70. Paley, E.L., Diet-Related Metabolic Perturbations of Gut Microbial Shikimate Pathway-Tryptamine-tRNA Aminoacylation-Protein Synthesis in Human Health and Disease. Int J Tryptophan Res, 2019. 12: p. 1178646919834550.

71. Binetti, J., et al., Deregulated Serotonin Pathway in Women with Morbid Obesity and NAFLD. Life (Basel), 2020. 10(10).

72. Hodge, S., et al., Obesity, whole blood serotonin and sex differences in healthy volunteers. Obes Facts, 2012. 5(3): p. 399–407.

73. Breisch, S.T., F.P. Zemlan, and B.G. Hoebel, Hyperphagia and obesity following serotonin depletion by intraventricular p-chlorophenylalanine. Science, 1976. 192(4237): p. 382–5.

74. Nonogaki, K., et al., Leptin-independent hyperphagia and type 2 diabetes in mice with a mutated serotonin 5-HT2C receptor gene. Nat Med, 1998. 4(10): p. 1152–6.

75. Ohara-Imaizumi, M., et al., Serotonin regulates glucose-stimulated insulin secretion from pancreatic β cells during pregnancy. Proc Natl Acad Sci U S A, 2013. 110(48): p. 19420–5.

76. Cirulli, E.T., et al., Profound Perturbation of the Metabolome in Obesity Is Associated with Health Risk. Cell Metab, 2019. 29(2): p. 488–500.e2.

77. Weir, J.M., et al., Plasma lipid profiling in a large population-based cohort. J Lipid Res, 2013. 54(10): p. 2898–908.

78. Klein, A.B., et al., Blood BDNF concentrations reflect brain-tissue BDNF levels across species. Int J Neuropsychopharmacol, 2011. 14(3): p. 347–53.

79. Pu, J., et al., Metabolomic changes in animal models of depression: a systematic analysis. Molecular Psychiatry, 2021. 26(12): p. 7328–7336.

80. Tavazzi, B., et al., Simultaneous high performance liquid chromatographic separation of purines, pyrimidines, N-acetylated amino acids, and dicarboxylic acids for the chemical diagnosis of inborn errors of metabolism. Clin Biochem, 2005. 38(11): p. 997–1008.

81. Kaul, R., et al., Canavan disease: mutations among Jewish and non-Jewish patients. Am J Hum Genet, 1994. 55(1): p. 34–41.

82. Chauhan, B.K., et al., Identification of differentially expressed genes in mouse Pax6 heterozygous lenses. Invest Ophthalmol Vis Sci, 2002. 43(6): p. 1884–90.

83. Parikh, S., et al., A clinical approach to the diagnosis of patients with leukodystrophies and genetic leukoencephelopathies. Mol Genet Metab, 2015. 114(4): p. 501–515.

84. Benson, M.D., et al., Severe retinal degeneration in a patient with Canavan disease. Ophthalmic Genet, 2021. 42(1): p. 75–78.

85. Narahara, M., et al., Immunohistochemical and chemical changes of beta-citryl-L-glutamate in the differentiation of bovine lens epithelial cells into lens fiber cells. Biol Pharm Bull, 2000. 23(6): p. 704–7.

86. Tsumori, M., et al., Presence of beta-citryl-L-glutamic acid in the lens: its possible role in the differentiation of lens epithelial cells into fiber cells. Exp Eye Res, 1995. 61(4): p. 403–11.

87. Hamada-Kanazawa, M., et al., *beta-Citryl-L-glutamate is an endogenous iron chelator that occurs naturally in the developing brain*. Biol Pharm Bull, 2010. 33(5): p. 729–37.

88. Krätzner, R., et al., A peroxisome proliferator-activated receptor gamma-retinoid X receptor heterodimer physically interacts with the transcriptional activator PAX6 to inhibit glucagon gene transcription. Mol Pharmacol, 2008. 73(2): p. 509–17.

89. Schinner, S., et al., Repression of glucagon gene transcription by peroxisome proliferator-activated receptor gamma through inhibition of Pax6 transcriptional activity. J Biol Chem, 2002. 277(3): p. 1941–8.

90. Zhu, J., et al., The role of pparγ in embryonic development of Xenopus tropicalis under triphenyltin-induced teratogenicity. Sci Total Environ, 2018. 633: p. 1245–1252.

91. Latta, L., et al., Abnormal neovascular and proliferative conjunctival phenotype in limbal stem cell deficiency is associated with altered microRNA and gene expression modulated by PAX6 mutational status in congenital aniridia. Ocul Surf, 2020.

92. Roux, L.N., et al., Modeling of Aniridia-Related Keratopathy by CRISPR/Cas9 Genome Editing of Human Limbal Epithelial Cells and Rescue by Recombinant PAX6 Protein. Stem Cells, 2018. 36(9): p. 1421–1429.

93. Lima Cunha, D., et al., Restoration of functional PAX6 in aniridia patient iPSC-derived ocular tissue models using repurposed nonsense suppression drugs. Mol Ther Nucleic Acids, 2023. 33: p. 240–253.

94. Zhao, P., et al., The TBK1/IKKε inhibitor amlexanox improves dyslipidemia and prevents atherosclerosis. JCI Insight, 2022. 7(17).

95. Oral, E.A., et al., Inhibition of IKKɛ and TBK1 Improves Glucose Control in a Subset of Patients with Type 2 Diabetes. Cell Metab, 2017. 26(1): p. 157–170.e7.

96. Reilly, S.M., et al., FGF21 is required for the metabolic benefits of IKKε/TBK1 inhibition. J Clin Invest, 2021. 131(10).

