## Supplementary data for "Metabolic and neuroactivity imbalances in plasma from aniridia patients with *PAX6* haploinsufficiency"

SUPPLEMENTARY RESULTS:

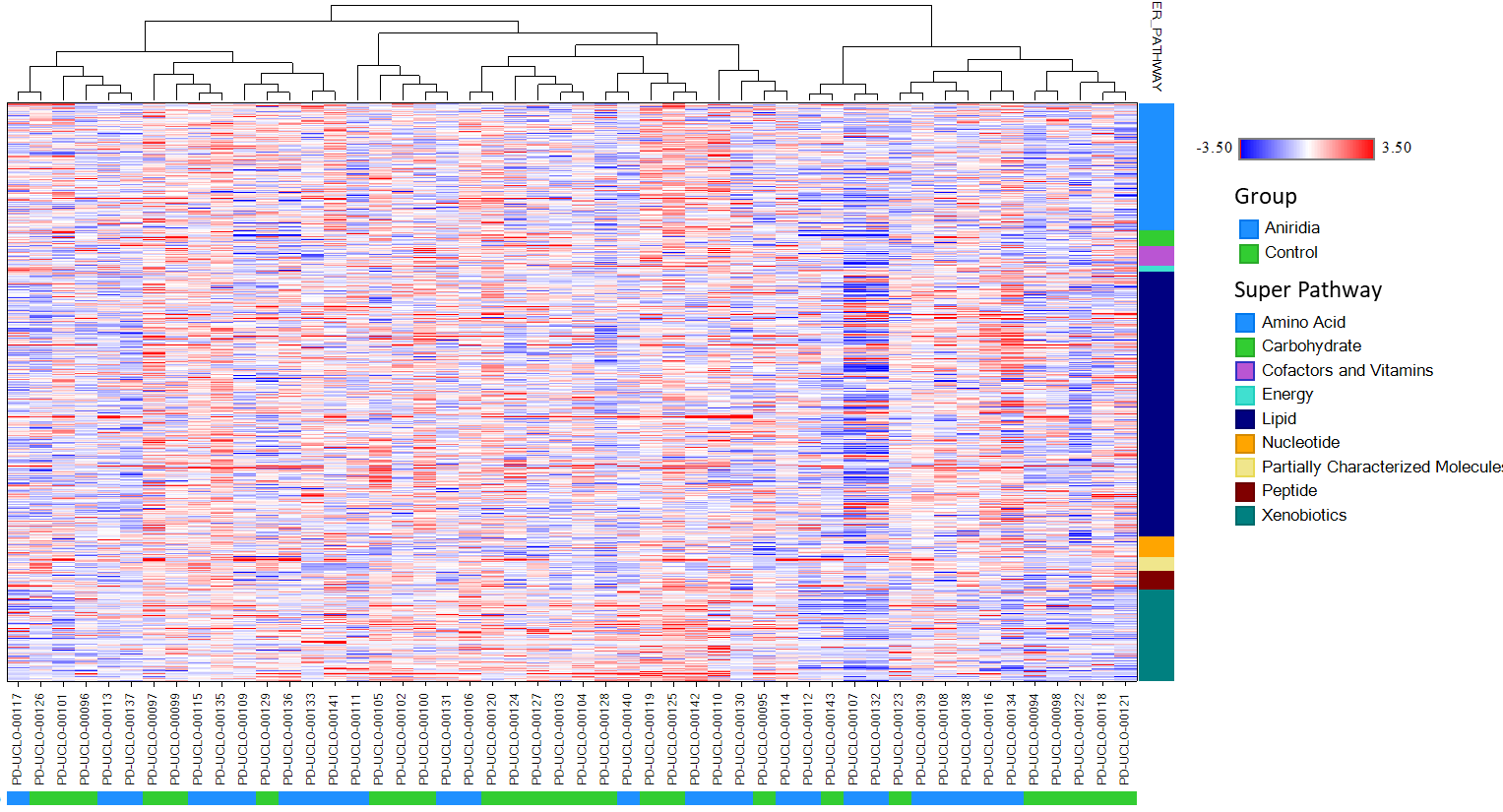

**Supplementary Figure 1.** Global heatmap of untargeted metabolomics dataset using hierarchical clustering (performed by Metabolon, Inc). Colours represent the fold change of each metabolite compared to mean control levels. Individual samples are distributed along the horizontal axis, with patient group samples highlighted in blue and control group in green. Compounds are shown along the vertical axis, grouped by super pathways represented in different colours.

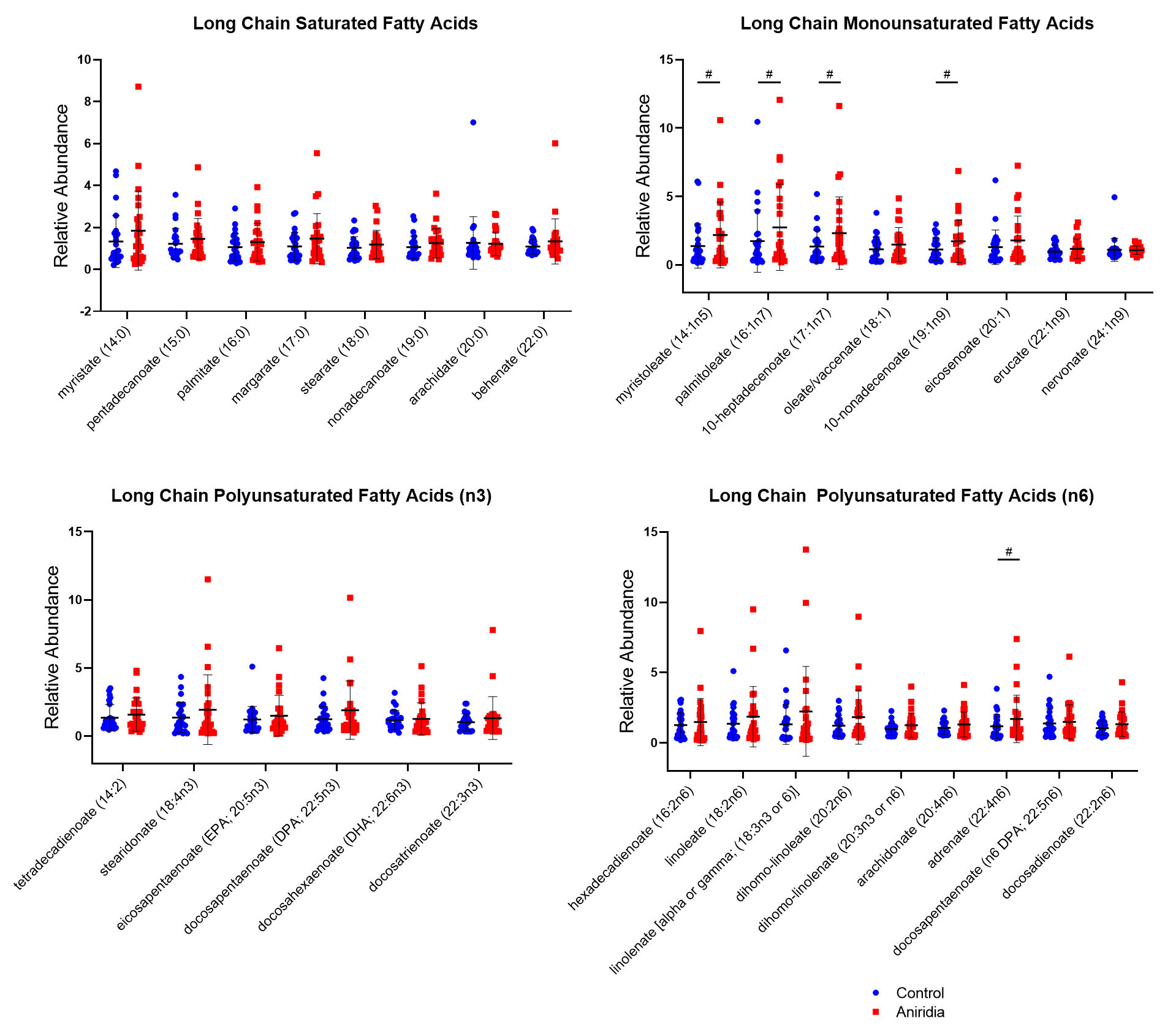
 **Supplementary Figure 2.** **Fatty acid metabolism alterations in aniridia patients plasma (*cont. from Fig 2*).** Relative abundance (in arbitrary units) of long-chain fatty acids, including saturated, monounsaturated and polyunsaturated (n3 and n6) in control (blue circles) and aniridia (red squares) groups. n=25 per group; ^#^ 0.05< p < 0.1.

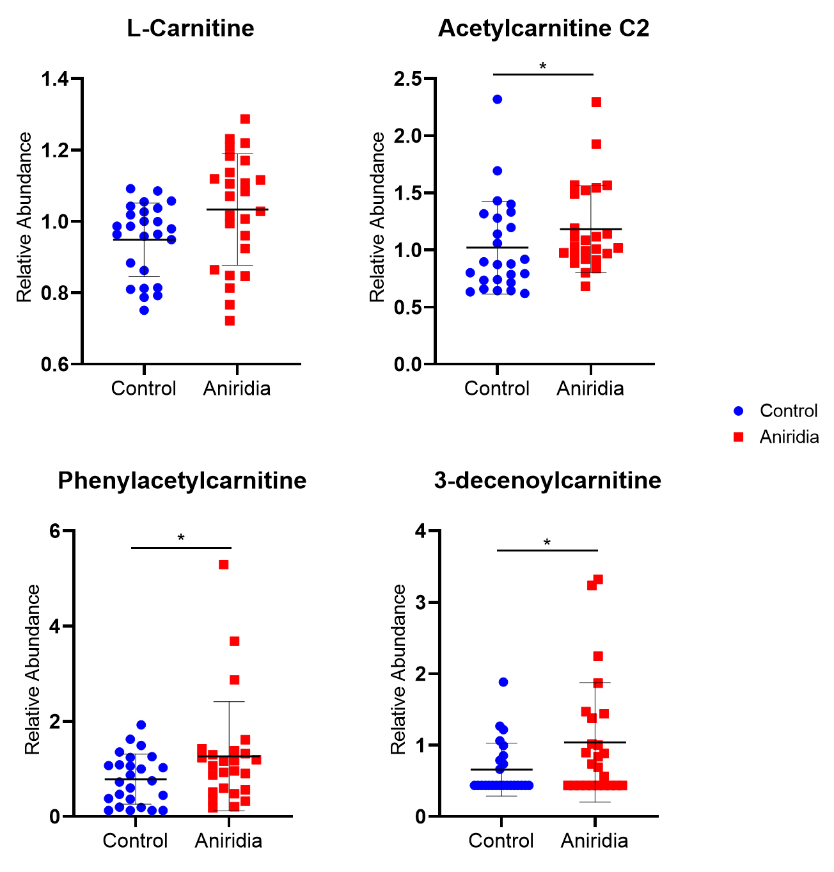

**Supplementary Figure 3**. Box plots showing the relative abundance of several carnitines detected in control (blue circles) and aniridia (red squares) groups. Black bars represent the mean and SD. n=25 pr group; * p ≤ 0.05.

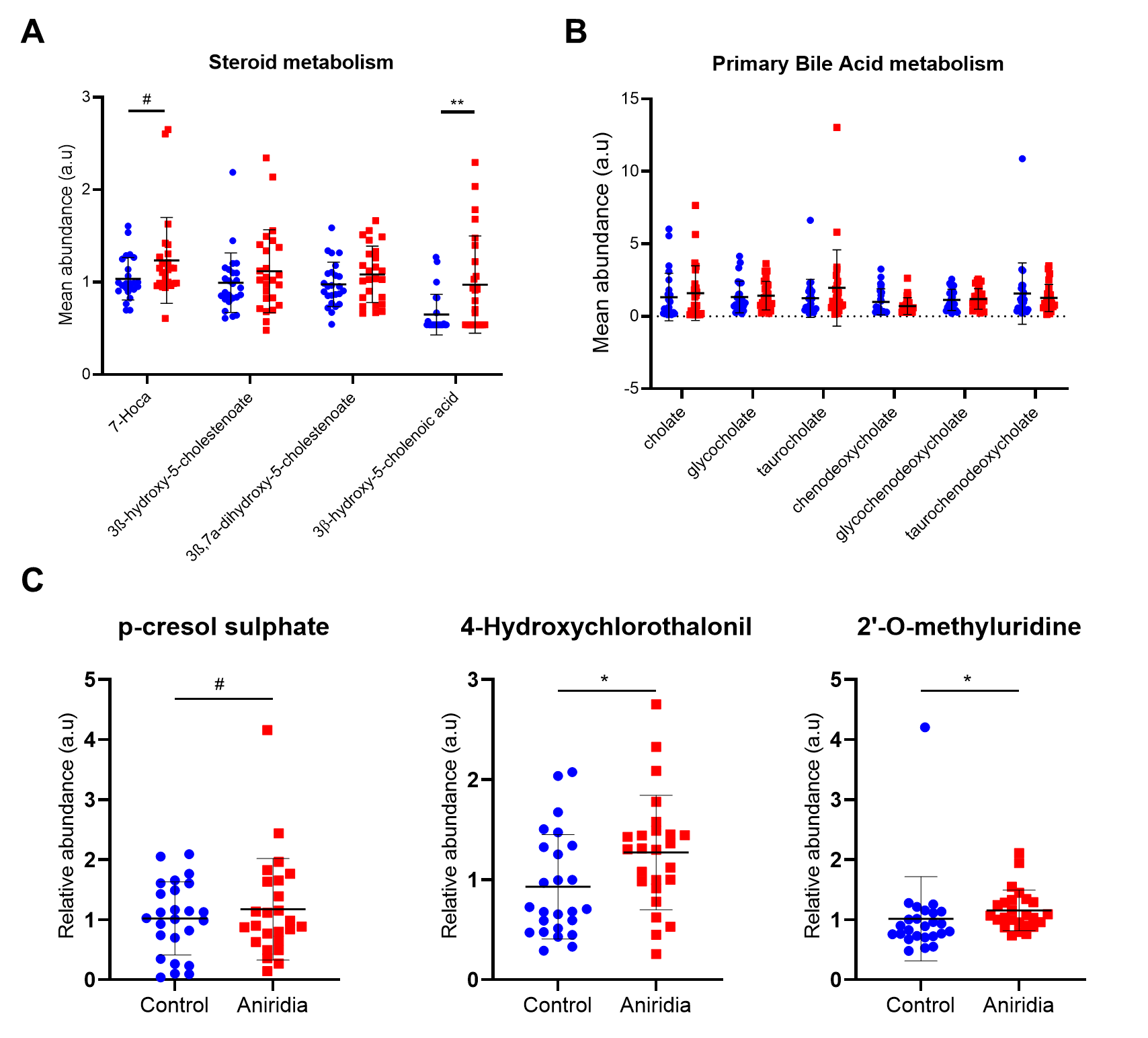

**Supplementary Figure 4.** Box plots showing the relative abundance of several metabolites from steroid metabolism (A), primary bile acid metabolism (B) and linked to gut microbiome (C). Control and aniridia groups values are shown in blue circles and red squares, respectively. Black bars represent the mean and SD. n=25 pr group; * p ≤ 0.05. Abbreviations: a.u. arbitrary units

SUPPLEMENTARY MATERIAL AND METHODS:

***Metabolon Platform***

**Sample Accessioning:** Following receipt, samples were inventoried and immediately stored at -80oC. Each sample received was accessioned into the Metabolon LIMS system and was assigned by the LIMS a unique identifier that was associated with the original source identifier only. This identifier was used to track all sample handling, tasks, results, etc. The samples (and all derived aliquots) were tracked by the LIMS system. All portions of any sample were automatically assigned their own unique identifiers by the LIMS when a new task was created; the relationship of these samples was also tracked. All samples were maintained at -80oC until processed.

**Sample Preparation:** Samples were prepared using the automated MicroLab STAR® system from Hamilton Company. Several recovery standards were added prior to the first step in the extraction process for QC purposes. To remove protein, dissociate small molecules bound to protein or trapped in the precipitated protein matrix, and to recover chemically diverse metabolites, proteins were precipitated with methanol under vigorous shaking for 2 min (Glen Mills GenoGrinder 2000) followed by centrifugation. The resulting extract was divided into five fractions: two for analysis by two separate reverse phase (RP)/UPLC-MS/MS methods with positive ion mode electrospray ionization (ESI), one for analysis by RP/UPLC-MS/MS with negative ion mode ESI, one for analysis by HILIC/UPLC-MS/MS with negative ion mode ESI, and one sample was reserved for backup. Samples were placed briefly on a TurboVap® (Zymark) to remove the organic solvent. The sample extracts were stored overnight under nitrogen before preparation for analysis.

**QA/QC:** Several types of controls were analyzed in concert with the experimental samples: a pooled matrix sample generated by taking a small volume of each experimental sample (or alternatively, use of a pool of well-characterized human plasma) served as a technical replicate throughout the data set; extracted water samples served as process blanks; and a cocktail of QC standards that were carefully chosen not to interfere with the measurement of endogenous compounds were spiked into every analyzed sample, allowed instrument performance monitoring and aided chromatographic alignment. Tables 1 and 2 describe these QC samples and standards. Instrument variability was determined by calculating the median relative standard deviation (RSD) for the standards that were added to each sample prior to injection into the mass spectrometers. Overall process variability was determined by calculating the median RSD for all endogenous metabolites (i.e., non-instrument standards) present in 100% of the pooled matrix samples. Experimental samples were randomized across the platform run with QC samples spaced evenly among the injections.

**Ultrahigh Performance Liquid Chromatography-Tandem Mass Spectroscopy (UPLC-MS/MS):** All methods utilized a Waters ACQUITY ultra-performance liquid chromatography (UPLC) and a Thermo Scientific Q-Exactive high resolution/accurate mass spectrometer interfaced with a heated electrospray ionization (HESI-II) source and Orbitrap mass analyzer operated at 35,000 mass resolution. The sample extract was dried then reconstituted in solvents compatible to each of the four methods. Each reconstitution solvent contained a series of standards at fixed concentrations to ensure injection and chromatographic consistency. One aliquot was analyzed using acidic positive ion conditions, chromatographically optimized for more hydrophilic compounds. In this method, the extract was gradient eluted from a C18 column (Waters UPLC BEH C18-2.1x100 mm, 1.7 μm) using water and methanol, containing 0.05% perfluoropentanoic acid (PFPA) and 0.1% formic acid (FA). Another aliquot was also analyzed using acidic positive ion conditions, however it was chromatographically optimized for more hydrophobic compounds. In this method, the extract was gradient eluted from the same afore mentioned C18 column using methanol, acetonitrile, water, 0.05% PFPA and 0.01% FA and was operated at an overall higher organic content. Another aliquot was analyzed using basic negative ion optimized conditions using a separate dedicated C18 column. The basic extracts were gradient eluted from the column using methanol and water, however with 6.5mM Ammonium Bicarbonate at pH 8. The fourth aliquot was analyzed via negative ionization following elution from a HILIC column (Waters UPLC BEH Amide 2.1x150 mm, 1.7 μm) using a gradient consisting of water and acetonitrile with 10mM Ammonium Formate, pH 10.8. The MS analysis alternated between MS and data-dependent MSn scans using dynamic exclusion. The scan range varied slighted between methods but covered 70-1000 m/z. Raw data files are archived and extracted as described below.

**Bioinformatics:** The informatics system consisted of four major components, the Laboratory Information Management System (LIMS), the data extraction and peak-identification software, data processing tools for QC and compound identification, and a collection of information interpretation and visualization tools for use by data analysts. The hardware and software foundations for these informatics components were the LAN backbone, and a database server running Oracle 10.2.0.1 Enterprise Edition.

**LIMS:** The purpose of the Metabolon LIMS system was to enable fully auditable laboratory automation through a secure, easy to use, and highly specialized system. The scope of the Metabolon LIMS system encompasses sample accessioning, sample preparation and instrumental analysis and reporting and advanced data analysis. All of the subsequent software systems are grounded in the LIMS data structures. It has been modified to leverage and interface with the in-house information extraction and data visualization systems, as well as third party instrumentation and data analysis software.

**Data Extraction and Compound Identification:** Raw data was extracted, peak-identified and QC processed using Metabolon’s hardware and software. These systems are built on a web-service platform utilizing Microsoft’s .NET technologies, which run on high-performance application servers and fiber-channel storage arrays in clusters to provide active failover and load-balancing. Compounds were identified by comparison to library entries of purified standards or recurrent unknown entities. Metabolon maintains a library based on authenticated standards that contains the retention time/index (RI), mass to charge ratio (*m/z)*, and chromatographic data (including MS/MS spectral data) on all molecules present in the library. Furthermore, biochemical identifications are based on three criteria: retention index within a narrow RI window of the proposed identification, accurate mass match to the library +/- 10 ppm, and the MS/MS forward and reverse scores between the experimental data and authentic standards. The MS/MS scores are based on a comparison of the ions present in the experimental spectrum to the ions present in the library spectrum. While there may be similarities between these molecules based on one of these factors, the use of all three data points can be utilized to distinguish and differentiate biochemicals. More than 3300 commercially available purified standard compounds have been acquired and registered into LIMS for analysis on all platforms for determination of their analytical characteristics. Additional mass spectral entries have been created for structurally unnamed biochemicals, which have been identified by virtue of their recurrent nature (both chromatographic and mass spectral). These compounds have the potential to be identified by future acquisition of a matching purified standard or by classical structural analysis.

**Curation:** A variety of curation procedures were carried out to ensure that a high quality data set was made available for statistical analysis and data interpretation. The QC and curation processes were designed to ensure accurate and consistent identification of true chemical entities, and to remove those representing system artifacts, mis-assignments, and background noise. Metabolon data analysts use proprietary visualization and interpretation software to confirm the consistency of peak identification among the various samples. Library matches for each compound were checked for each sample and corrected if necessary.

**Metabolite Quantification and Data Normalization:** Peaks were quantified using area-under-the-curve. For studies spanning multiple days, a data normalization step was performed to correct variation resulting from instrument inter-day tuning differences. Essentially, each compound was corrected in run-day blocks by registering the medians to equal one (1.00) and normalizing each data point proportionately (termed the “block correction”). For studies that did not require more than one day of analysis, no normalization is necessary, other than for purposes of data visualization. In certain instances, biochemical data may have been normalized to an additional factor (e.g., cell counts, total protein as determined by Bradford assay, osmolality, etc.) to account for differences in metabolite levels due to differences in the amount of material present in each sample.

**Metabolic Pathway Networks and Analysis**: To visualise and analyse small molecules within relevant networks of metabolic pathways, the detected metabolites in Aniridia patient and healthy control study groups were subjected to MetaboLync pathway analysis (MPA) software (portal.metabolon.com). Significantly altered pathways were determined by completing pathway set enrichment analysis within MPA software which was determined by the following equation:

\# of significant metabolites in pathway (k)/ total # of detected metabolites in pathway (m)/ total # of significant metabolites (n)/ total # of detected metabolites (N) or (k/m)/(n/N).

A pathway impact score greater than 1 indicates that the pathway contains a higher number of experimentally regulated compounds relative to the overall study in Aniridia patients relative to healthy controls. Biological relevance of metabolites was assessed using the Human Metabolome Database (<https://hmdb.ca>).
