## Supplementary Figure 2 for "Metabolic and neuroactivity imbalances in plasma from aniridia patients with *PAX6* haploinsufficiency"

**Supplementary Table 2**: Comparison of nutrient intakes between aniridia cases and controls

| **Nutrient** |  | Aniridia cases (n=25) | |  | Controls (n=25) | |  | P-value* |
| --- | --- | --- | --- | --- | --- | --- | --- | --- |
|  |  | Mean | SD |  | Mean | SD |  |  |
| Carbohydrate (g) |  | 279.2 | 92.4 |  | 249.1 | 91.4 |  | 0.29 |
| Carbohydrate as a % of total energy |  | 49.7 | 4.4 |  | 47.7 | 5.3 |  | 0.12 |
| Total sugars (g) |  | 125.8 | 47.6 |  | 114.1 | 48.7 |  | 0.39 |
| Glucose (g) |  | 25.8 | 12.8 |  | 22.7 | 10.0 |  | 0.35 |
| Fructose (g) |  | 29.3 | 14.6 |  | 25.8 | 12.7 |  | 0.30 |
| Sucrose (g) |  | 47.1 | 20.0 |  | 45.6 | 26.6 |  | 0.53 |
| Maltose (g) |  | 4.72 | 2.48 |  | 3.65 | 1.70 |  | 0.16 |
| Lactose (g) |  | 17.5 | 10.6 |  | 15.0 | 10.8 |  | 0.40 |
| Starch (g) |  | 151.7 | 53.9 |  | 133.5 | 57.2 |  | 0.29 |
| Energy (kJ) |  | 8960 | 2776 |  | 8342 | 3002 |  | 0.43 |
| Total fat as a % of total energy |  | 33.6 | 4.1 |  | 34.7 | 3.9 |  | 0.26 |
| Saturated fat as a % of total energy |  | 12.4 | 2.5 |  | 12.6 | 2.4 |  | 0.79 |
| Monounsaturated fat as a % of total energy |  | 11.7 | 1.9 |  | 12.2 | 1.4 |  | 0.18 |
| PUFAs as a % of total energy |  | 6.62 | 1.59 |  | 6.95 | 2.13 |  | 0.88 |
| N3 PUFAs as a % of total energy |  | 0.71 | 0.23 |  | 0.76 | 0.23 |  | 0.21 |
| Protein as a % of total energy |  | 16.8 | 2.6 |  | 17.2 | 3.1 |  | 0.42 |
| Fibre (g) |  | 20.4 | 6.2 |  | 18.3 | 7.5 |  | 0.27 |
| Alcohol (g) |  | 3.71 | 9.05 |  | 5.14 | 10.19 |  | >0.99 |
| Thiamin (mg) |  | 1.94 | 0.51 |  | 1.64 | 0.59 |  | 0.10 |
| Riboflavin (mg) |  | 2.26 | 1.02 |  | 1.89 | 0.74 |  | 0.26 |
| Niacin (mg) |  | 22.5 | 7.4 |  | 21.2 | 7.8 |  | 0.58 |
| Vitamin B6 (mg) |  | 1.99 | 0.69 |  | 1.79 | 0.65 |  | 0.28 |
| Vitamin B12 (µg) |  | 7.78 | 7.37 |  | 5.78 | 2.94 |  | 0.40 |
| Folate (µg) |  | 356 | 124 |  | 293 | 134 |  | 0.10 |
| Carotene (µg) |  | 5086 | 2264 |  | 4062 | 2512 |  | 0.07 |

* Mann-Whitney test
